# Root microbiota assembly and adaptive differentiation among European *Arabidopsis* populations

**DOI:** 10.1101/640623

**Authors:** Thorsten Thiergart, Paloma Durán, Thomas Ellis, Ruben Garrido-Oter, Eric Kemen, Fabrice Roux, Carlos Alonso-Blanco, Jon Ågren, Paul Schulze-Lefert, Stéphane Hacquard

## Abstract

Factors that drive continental-scale variation in root microbiota and plant adaptation are poorly understood. We monitored root-associated microbial communities in *Arabidopsis thaliana* and co-occurring grasses at 17 European sites across three years. Analysis of 5,625 microbial community profiles demonstrated strong geographic structuring of the soil biome, but not of the root microbiota. Remarkable similarity in bacterial community composition in roots of *A. thaliana* and grasses was explained by the presence of a few diverse and geographically widespread taxa that disproportionately colonize roots across sites. In a reciprocal transplant between two *A. thaliana* populations in Sweden and Italy, we uncoupled soil from location effects and tested their respective contributions to root microbiota variation and plant adaptation. The composition of the root microbiota was affected by location and soil origin, and to a lesser degree by host genotype. The filamentous eukaryotes were particularly strongly affected by location. Strong local adaptation between the two *A. thaliana* populations was observed, with difference in soil properties and microbes of little importance for the observed magnitude of adaptive differentiation. Our results suggest that, across large spatial scales, climate is more important than are soil conditions for plant adaptation and variation in root-associated filamentous eukaryotic communities.

## Introduction

Plants interact with multi-kingdom microbial communities (e.g. bacteria, fungi, oomycetes) that can impact host fitness, either directly, or indirectly through microbe-microbe interactions^1, 2, 3, 4, 5^. The immune system of plants, rhizodeposits, and microbial interactions are known determinants of root-associated microbial assemblages and make them distinct from the surrounding soil biota^6, 7, 8, 3^. Large-scale spatial variation in the composition of the soil biota has been associated with difference in edaphic and climatic conditions^9^. Particularly, local edaphic factors such as soil pH primarily predict geographic distribution of soil bacteria^10, 11, 12^, whereas climatic variables better predict fungal distribution in soil^13^. However, systematic field studies exploring and disentangling the extent to which variation in soil and climatic conditions impacts root microbiota composition and adaptive differentiation in plants are lacking.

Local adaptation has been documented in a large number of plant species and across both small and large spatial scales^14^. For example, reciprocal transplants and common-garden experiments have provided evidence of strong adaptive differentiation among natural populations of the model plant *Arabidopsis thaliana*^15, 16^. However, the relative importance of different abiotic and biotic factors for the evolution and maintenance of local adaptation is poorly known^17, 18^. Particularly, soil edaphic factors and soil microbes are known to influence flowering phenology and modulate host fitness in natural soils^19, 20, 21, 22, 23^, even at the scale of a few meters^24^. Yet, information about the extent to which differences in soil properties contribute to divergent selection and the maintenance of adaptive differentiation among plant populations is still limited beyond classical examples of adaptation to extreme soil conditions^25^.

Here, we tested whether roots of *A. thaliana* and co-occurring grasses growing in various soils and climatic environments establish stable associations with bacterial and filamentous eukaryotic communities across a latitudinal gradient in Europe. In a reciprocal transplant between two *A. thaliana* populations in Sweden and Italy, we uncoupled soil from location effects and experimentally tested the hypothesis that soil properties and climate drive root microbiota assembly and adaptive differentiation between the two *A. thaliana* populations.

We found that a widespread set of bacteria, but not filamentous eukaryotes, establish stable associations with roots of *A. thaliana* and grasses across 17 sites in Europe, despite strong geographical structuring and variation in the surrounding soil communities. The reciprocal transplant of soil and plant genotypes between two native *A. thaliana* populations in northern and southern Europe showed that the composition of root microbiota was more affected by soil properties and location than by host genotype. The effect of soil was stronger than that of location for root-associated bacteria, whereas the effect of location was stronger for root-associated fungi and oomycetes. Transplant location, rather than origin of soil, also largely accounted for strong selection against the nonlocal *A. thaliana* genotype at each site. Our results suggest that climate is a primary force driving geographic variation in filamentous eukaryotic communities in roots and adaptive differentiation between *A. thaliana* populations in northern and southern Europe.

## Results

### Continental-scale survey of the *A. thaliana* root microbiota

We sampled natural *A. thaliana* populations at the flowering stage at 17 sites along a latitudinal gradient in Europe in three consecutive years (2015, 2016 and 2017). We harvested bulk soil (soil), rhizosphere (RS), rhizoplane (RP), and root endosphere (root) compartments of *A. thaliana* and co-occurring grasses (**Supplementary Fig. 1a, b**) at four sites in Sweden (SW1-4), six in Germany (GE1-6), three in France (FR1-3), one in Italy (IT1), and three in Spain (SP1-3) (**Fig. 1a**), each having distinct environmental and soil characteristics (**Supplementary Table 1**). DNA was isolated and microbial community composition analyzed for a total of 1,125 samples. Bacterial, fungal and oomycetal communities were profiled using primer pairs targeting the V2V4 and V5V7 regions of the bacterial 16S rRNA gene, the ITS1 and ITS2 segments of the fungal ITS, and the ITS1 segment of the oomycetal ITS, resulting in the sequencing of 5,625 microbial community profiles (**Supplementary Table 2**). Given the correlation observed at the class level between the independently used primer pairs that target the two regions of the bacterial 16S rRNA gene (Pearson correlation, r = 0.40, p < 0.001) and the two regions of the fungal ITS (Pearson correlation, r = 0.93, p < 0.001) (**Supplementary Fig. 1c**), only the bacterial V5V7 and fungal ITS1 variable segments were considered for later analyses.

**Fig 1:**
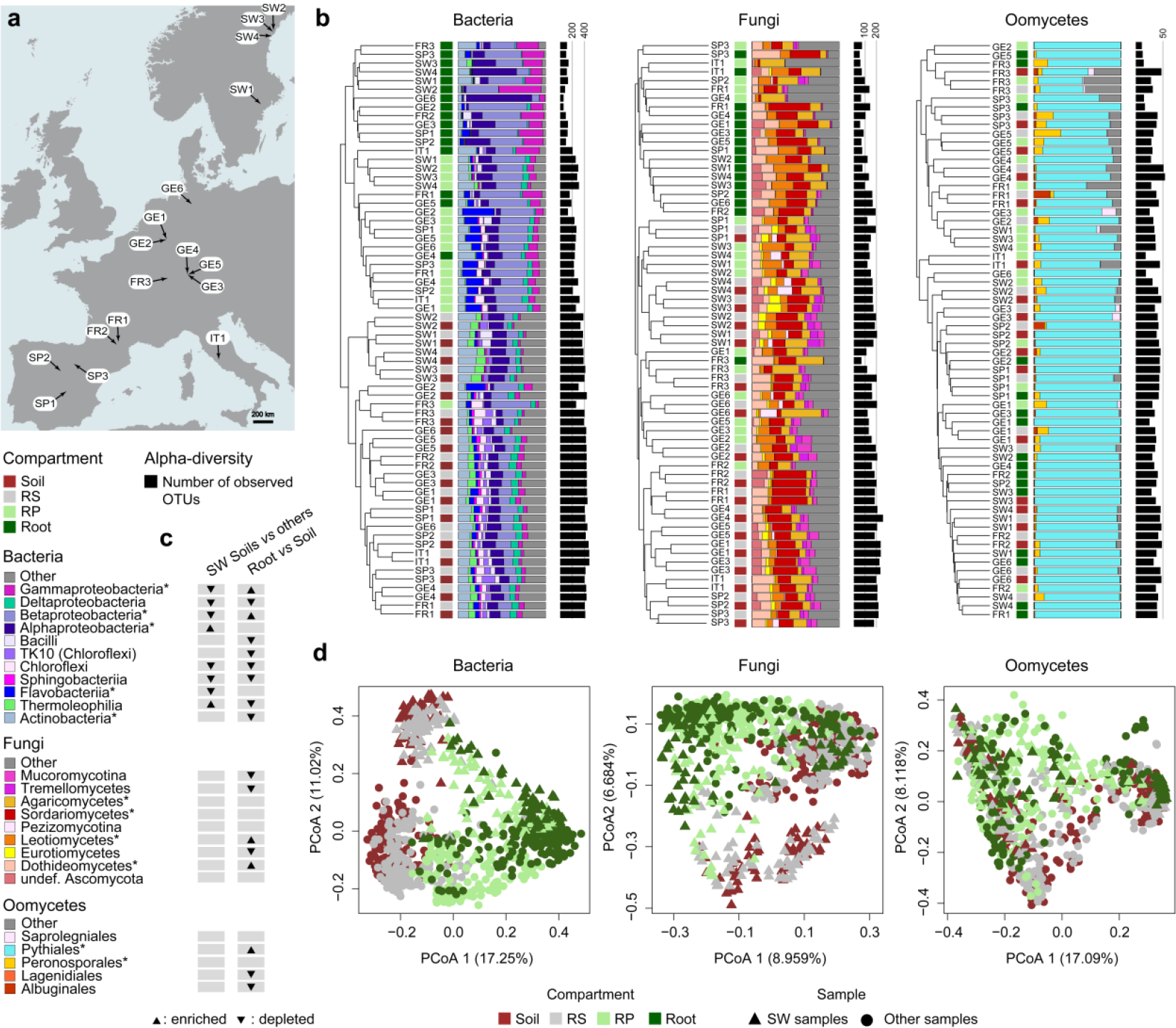
Microbial community structure in 17 European *A. thaliana* populations. **a**, European map showing names and locations of the 17 *A. thaliana* populations. **b**, Bray-Curtis similarity-based dendrogram showing averaged bacterial (left), fungal (middle), oomycetal (right) community composition for each compartment at each site. Only OTUs with relative abundance > 0.1% were considered. The total number of processed samples was 896 and only those with more than 1,000 reads were used to calculate average Bray-Curtis distances. Compartments are indicated with colored squares: soil (dark red), rhizosphere (RS, grey), rhizoplane (RP, light green), root (dark green). For each sample, community composition (class or order level) is indicated with bar plots and microbial alpha-diversity is represented with black bars according to the number of observed OTUs in the corresponding rarefied datasets (1,000 reads). **c**, Differential abundance analysis (class or order level) between the four Swedish soil samples and the other 13 soils, as well as between root and soil samples. Triangles depict statistically significant differences (Wilcoxon rank sum test, FDR < 0.05). **d**, Principal coordinate analysis (PCoA) based on Bray Curtis distances between samples harvested across 17 sites, four compartments and three successive years (2015, 2016, 2017). Microbial communities are presented for the whole *A. thaliana* dataset for bacteria (n = 881), fungi (n = 893), and oomycetes (n = 875) and color-coded according to the compartment. RS: rhizosphere. RP: rhizoplane. Triangles represent the Swedish samples and circles all the other samples. OTUs with relative abundance < 0.1% were excluded from the dataset.

### Convergence in root microbiota composition across European *A. thaliana* populations

If plant roots establish stable associations with microbial communities across large geographical distances, we expect a strong host filtering effect on root microbiota composition. Inspection of alpha-diversity indices (i.e. Shannon index, number of observed OTUs) revealed a gradual decrease of bacterial, fungal, and oomycetal diversity from the soil to the root endosphere (Kruskal-Wallis test, p < 0.05), with significantly stronger decrease for root-associated bacteria than for filamentous eukaryotes (**Fig. 1b** and **Supplementary Fig. 2a, b**). Analysis of microbial community structure based on Bray-Curtis distances across sites and compartments revealed that bacterial communities in the root endosphere and RP cluster by compartments, and by site in the RS and soil (**Fig. 1b**). A clustering by compartment was also visible among root endosphere samples for fungi, but not for oomycetes (**Fig. 1b**). By considering soil and root endosphere samples, compartment explained variation in bacterial community composition more than did site (Compartment: 18.9%; site: 17.2%, PERMANOVA with Bray–Curtis distances, p < 0.01; **Supplementary Table 3**). In contrast, site explained variation in fungal and oomycetal community composition more than did compartment (Site: 20.2% for fungi, 15.5% for oomycetes; Compartment: 6.5% for fungi, 2.6% or oomycetes; PERMANOVA with Bray–Curtis distances, p < 0.01, **Supplementary Table 3**). Nonetheless, the observation that endosphere-associated bacterial and fungal communities show overall more similarities across sites than across compartments at a given site implies structural convergence of the *A. thaliana* root microbiota at a continental scale. This is well illustrated by the marked differences in soil biota observed between Swedish soils (SW1-4) and the other European soils (**Fig.1c** and **d, Supplementary Fig. 2c**), which is largely diminished in the corresponding root endosphere samples (**Fig. 1d**). Compared to surrounding soil samples, microbial communities in plant roots showed a significant enrichment of taxa belonging to the bacterial classes Beta- and Gamma-Proteobacteria, the fungal classes Leotiomycetes and Dothideomycetes and the oomycetal order Pythiales (Wilcoxon rank sum test, FDR < 0.05, **Fig. 1c** and **Supplementary Fig. 2d**). Our results demonstrate that the root environment drives remarkable convergence in bacterial, and to a lesser extent fungal and oomycetal community composition across European sites separated by up to 3,500 km, despite considerable differences in soil properties (**Supplementary Table 1**) and soil microbial communities.

**Fig 2:**
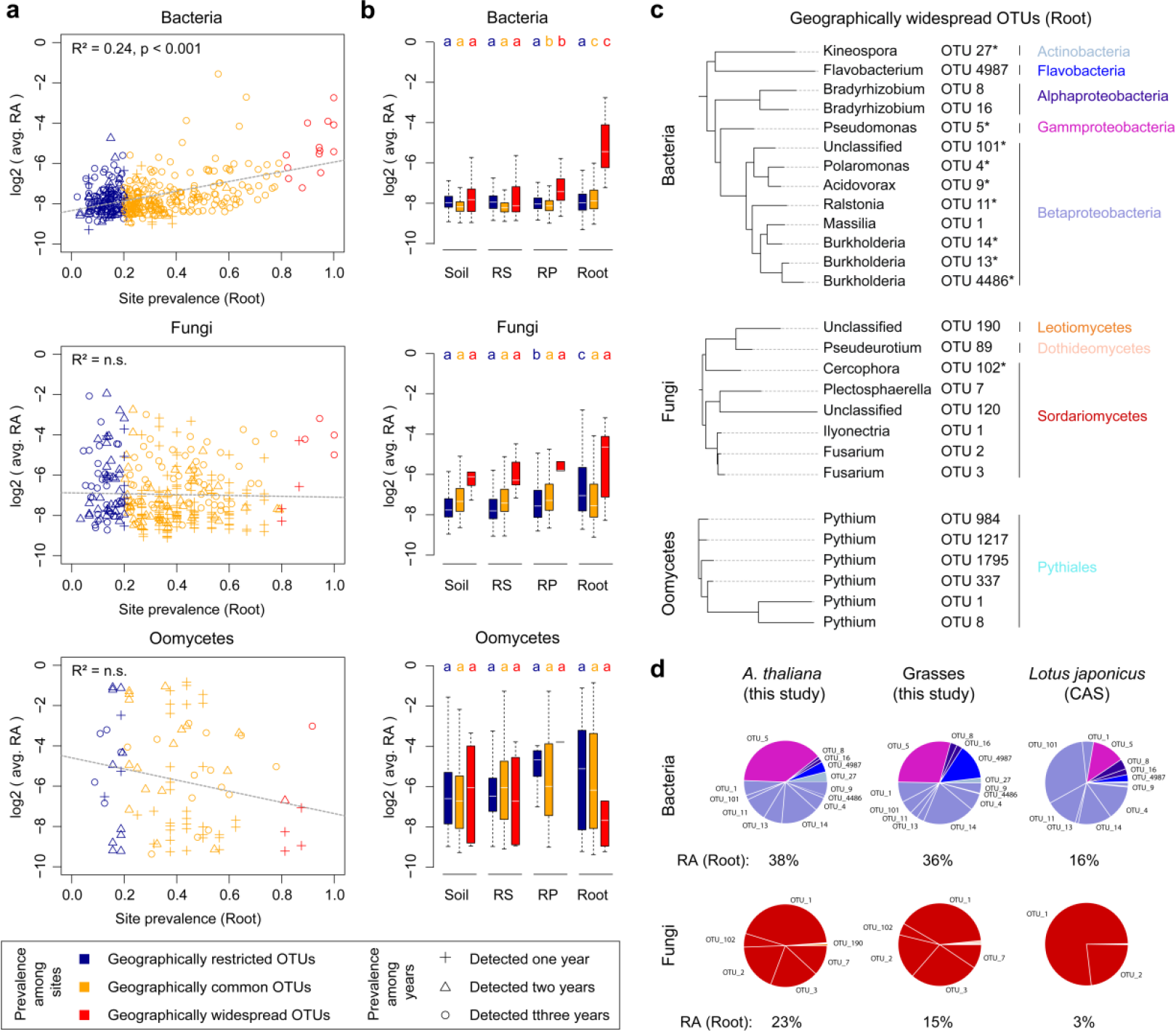
Geographically widespread taxa in the roots of *A. thaliana* and grasses. **a**, Correlation between OTUs prevalence across sites in plant roots and averaged OTUs relative abundance (RA, log2) in plant roots. Bacteria: upper panel. Fungi: middle panel. Oomycetes: lower panel. For calculating averaged RA, only samples where the actual OTUs are present were considered. Blue: geographically restricted OTUs (site prevalence < 20%). Orange: geographically common OTUs (site prevalence 20-80%). Red: geographically widespread OTUs (site prevalence > 80%). The different shapes highlight root-associated OTUs detected one year, or across two or three years. RA and prevalence are averaged across the years where one OTU is present. **b**, Boxplots of the averaged RA (log2) of geographically restricted, common and widespread OTUs detected in each of the four compartments. Letters depict significant differences across compartments (Kruskal-Wallis test, FDR < 0.01). **c**, Phylogenetic trees of geographically widespread root-associated OTUs (red symbols in panel a) constructed based on the 16s rRNA V5V7 gene fragments (bacteria) and the ITS1 region (fungi, oomycetes). Microbial OTUs significantly enriched in root compared to soil samples are indicated with a star (FDR < 0.05) **d**, Geographically widespread OTUs detected in roots of *A. thaliana* and conserved signatures in roots of grasses and *Lotus japonicus.* The RA and proportion of widespread bacterial and fungal OTUs detected in *A. thaliana* roots are shown for *A. thaliana* (17 sites), co-occuring grasses (17 sites), as well as for *Lotus japonicus* grown in the Cologne Agricultutral Soil (CAS). All shown OTUs have RA > 0.1%. The total RA of these OTUs in root samples is indicated below the circular diagrams.

### Root endosphere bacteria and *A. thaliana* exhibit stable associations across Europe

Given the limited variation in microbial community composition observed in plant roots across geographically distant sites, we hypothesized that the presence of geographically widespread microbes might contribute to convergence in root microbiota composition. To identify microbial OTUs that are widely distributed in roots of *A. thaliana* across Europe, we calculated their prevalence across all 17 sites and those detected in more than 80% of the sites were defined as geographically widespread (**Fig. 2a** and **Supplementary Fig. 3a**). Remarkably, we observed a positive correlation (linear regression, R^2^ = 0.24, p < 0.001) between the prevalence of root-associated bacteria across sites and their relative abundance (RA) in root endosphere samples, suggesting that bacterial taxa that colonize *A. thaliana* roots across Europe are also the most abundant in this niche (**Fig. 2a**). This implies that a small subset of bacterial taxa have evolved mechanisms to dominate the bacterial root microbiota at a continental scale, irrespective of major differences in the surrounding local bacterial soil biota. In contrast, no significant correlation was observed between relative abundance in plant roots and prevalence across sites for root-associated fungi and oomycetes (linear regression, p > 0.05, **Fig. 2a**). By inspecting the abundance profiles of OTUs with restricted or widespread geographic distribution across compartments (**Fig. 2b**), we observed that geographically widespread bacterial OTUs are significantly more abundant in RP and Root samples than in the corresponding soil samples (Kruskal-Wallis test, FDR < 0.01). In contrast, fungal OTUs that have a narrow geographical distribution are significantly more abundant in RP and Root samples than in the corresponding bulk soil (Kruskal-Wallis test, FDR < 0.01).

**Fig 3:**
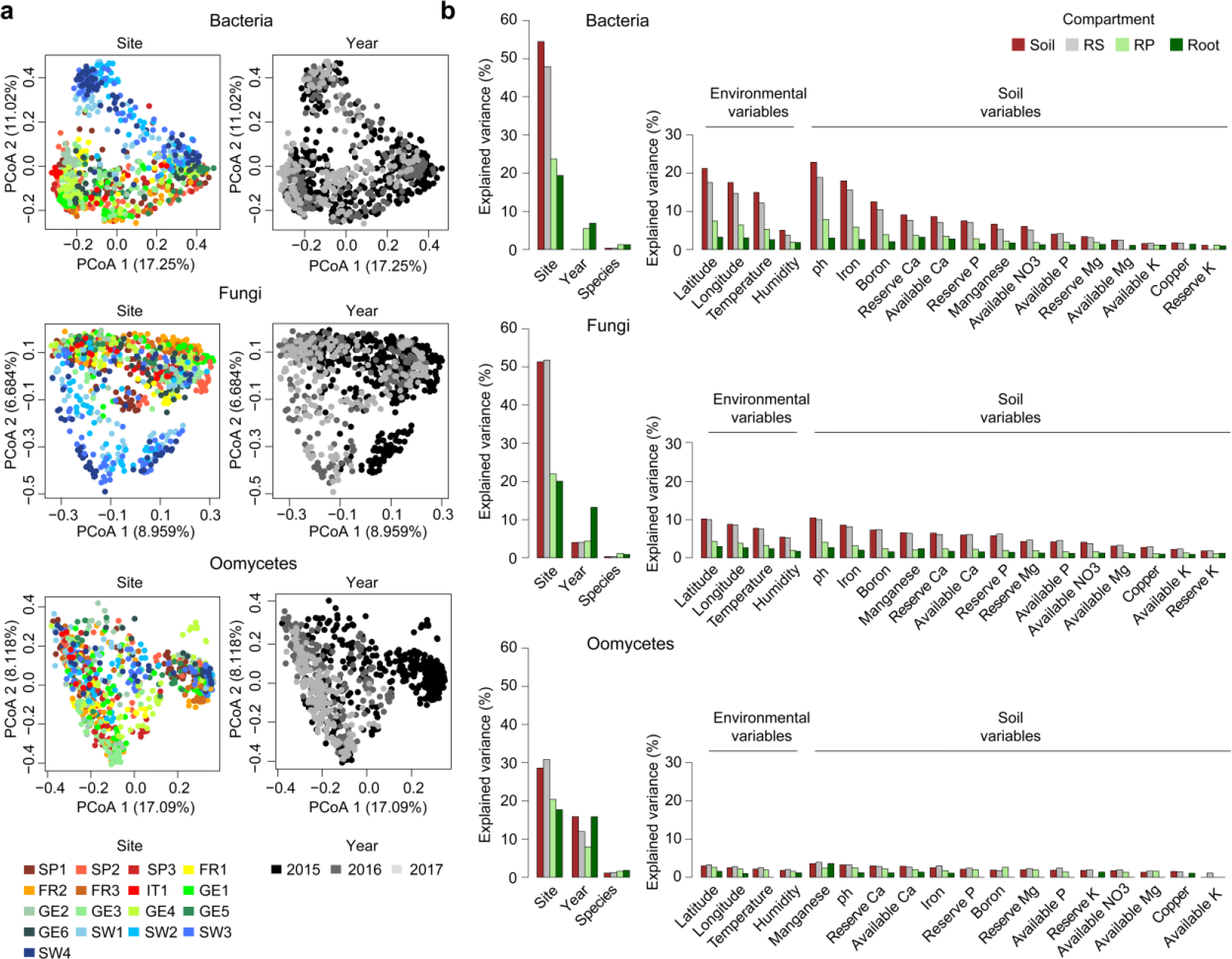
Factors shaping the *A. thaliana* root microbiota at a continental scale. **a**, Principal coordinate analysis (PCoA) based on Bray Curtis distances between samples harvested across 17 sites, four compartments and three successive years (2015, 2016, 2017). Microbial communities are presented for the whole *A. thaliana* dataset for bacteria (upper panel, 881 samples), fungi (middle panel, 893 samples), and oomycetes (lower panel, 875) and color-coded either according to the site or the harvesting year. OTUs with relative abundance < 0.1% were excluded from the dataset. **b**, Effect of site, harvesting year, host species, as well as of individual soil and environmental variables measured at each site, on bacterial, fungal, and oomycetal community composition. Explained variance (%) for each explanatory variable is shown for the different compartments and ranked according to the best explanatory variable in soil. The percentage of explained variance for each parameter was calculated based on permutational multivariate analysis of variance and only significant associations are shown (PERMANOVA, p < 0.01). RS: rhizosphere. RP: rhizoplane.

We identified 13 geographically widespread bacterial OTUs that are consistently detected in the root endosphere of *A. thaliana* across European sites (**Fig. 2a, c** and **Supplementary Table 4**), accounting for 38% of the total relative abundance in this niche (**Fig. 2d**). These taxa belong to 5 different classes, and cover 10 bacterial genera, including *Bradyrhizobium, Pseudomonas, Polaromonas, Acidovorax, Ralstonia, Massilia, Burkholderia, Kineospora* and *Flavobacterium* (**Fig. 2c**), which indicates convergent adaptation to the root environment in phylogenetically distant bacterial lineages at a continental scale. Notably, these 13 bacterial OTUs were detected across all three years (**Fig. 2a**), and nine of them were significantly enriched in plant roots compared to soil (FDR < 0.05, highlighted with a star in **Fig. 2c**). In contrast, the abundance of the 14 geographically widespread OTUs of root-associated filamentous eukaryotes varied among years, and were dominated by fungi from only three classes (Sordariomycetes, Leotiomycetes and Dothideomycetes) and oomycetes from a single genus (*Pythium*), and were not particularly root-enriched (**Fig. 2a, c, Supplementary Table 4**). Our results suggest that the few geographically widespread bacteria that abundantly colonize roots of *A. thaliana* drive convergence in bacterial community composition at a continental scale.

### Geographically widespread bacterial OTUs in *A. thaliana* roots are ubiquitous in distantly related plant species

The conserved taxa that consistently associate with roots of *A. thaliana* might represent a widespread microbial multi-kingdom community that also associate with distantly related plant species. To test this hypothesis, we harvested co-occurring grasses at each of the 17 sites across Europe and compared microbial community composition between roots of *A. thaliana* and grasses. The factor host species weakly, but significantly explained bacterial, fungal, and oomycetal community composition in root endosphere samples when considering the whole dataset (1.3%, 0.89% 1.8% of the variance, respectively, PERMANOVA with Bray–Curtis distances, p < 0.01, **Supplementary Table 5**). The overall effect of host species is likely underestimated because we could not sample the same grass species at all sites and we pooled plant individuals for efficient RP and root fractionation (see **Methods**). By inspecting the species effect at each site separately using PERMANOVA, we observed a stronger effect of the host species on the root microbiota, although these differences were significant for only few sites (average explained variance: 11% for bacteria, 8.8% for fungi, 7.7% for oomycetes, **Supplementary Table 6**). Comparison of microbial OTU prevalence in roots of *A. thaliana* with those of co-occurring grasses revealed overall consistency in root OTU prevalence at a continental scale (Spearman rank correlation, r_s_ = 0.69 for bacteria; r_s_ = 0.79 for fungi; r_s_ = 0.72 for oomycetes; p *<* 0.01] (**Supplementary Fig. 3b**). This indicates an overall conserved distribution of geographically restricted and widespread OTUs in roots of phylogenetically distant plants species that evolved independently in the Brassicaceae and Poaceae lineages. Inspection of the 13 geographically widespread bacterial OTUs detected in roots of *A. thaliana* revealed that these are also abundantly detected in roots of co-occurring grasses (13/13 detected, 36% of the total relative abundance), whereas conservation was less obvious for the geographically widespread fungal OTUs (5/7 detected, 15% of the total relative abundance). Further inspection of their abundance in roots of *Lotus japonicus* (Fabaceae) grown in a completely different soil type (i.e. Cologne Agricultural Soil^26^) validated the ubiquitous nature of the bacterial OTUs (11/13 detected, 16% of the total RA in roots), but not of the fungal OTUs (2/8 detected, 3% of the total RA in roots; **Fig. 2d**). The results suggest that a small number of geographically widespread bacteria can efficiently colonize roots of distantly related plant species and establish potentially stable associations with plant roots over evolutionary time.

### Spatial and temporal variation in root microbiota differentiation

Despite limited among-site variation in microbial communities in root endosphere samples across European sites, we did observe spatial and temporal variation in their composition (**Fig. 3a, Supplementary Fig. 4a, b**). Site explained 19.4%, 20.1% and 17.7% of the variance in bacterial, fungal, and oomycetal community composition in roots of European *A. thaliana* populations respectively, compared with 54.5%, 51.3%, and 28.6% of the variance in corresponding soil samples (PERMANOVA with Bray–Curtis distances, p < 0.01, **Fig. 3b** and **Supplementary Table 5**). Inspection of correlations between environmental or soil variables and microbial community composition revealed a gradual decrease in explanatory power from soil towards root compartments, as well as stronger correlations for bacterial communities than for filamentous eukaryotes (**Fig. 3b, Supplementary Fig. 5**, and **Supplementary Tables 1** and **5**). Although part of the observed correlations can be due to confounding effects between variables (**Supplementary Fig. 5a, b**), latitude and soil pH explained the highest proportions of variation in bacterial and fungal community composition in soil (bacteria: 21.2% and 22.8%, respectively; fungi: 10.2% and 10.5%, respectively; PERMANOVA, p < 0.01), and remained among the most significant variables explaining among-site variation in the composition of the root microbiota across European *A. thaliana* populations (bacteria: 3.2% and 3.0%, respectively; fungi: 3.0% and 2.7%, respectively; PERMANOVA, p < 0.01) (**Fig. 3b, Supplementary Fig. 5** and **Supplementary Table 5**). To assess among-year variation in root microbiota composition, we sampled *A. thaliana* and grass populations at the same phenological stage in three successive years in spring 2015, 2016, and 2017. Remarkably, “year” explained more variation in root-associated microbial communities (bacteria: 6.9%, fungi: 13.2%, oomycetes: 15.8%, PERMANOVA, p < 0.01), than did “host species” (< 2%, PERMANOVA, p < 0.01), suggesting that year-to-year environmental variation affected the establishment of the root microbiota more than did differences between hosts separated by > 140 million years of reproductive isolation^27^ (**Fig. 3b** and **Supplementary Fig. 4b** and **Supplementary Table 5**). Among-year variation was particularly strong for soil- and root-associated fungal (soil: 4.0 %, root: 13.2%, PERMANOVA, p < 0.01) and oomycetal (soil: 15.9%, root: 15.8%, PERMANOVA, p < 0.01) communities (**Fig. 3a, b**, and **Supplementary Fig. 4b** and **Supplementary Table 5**).

**Fig 4:**
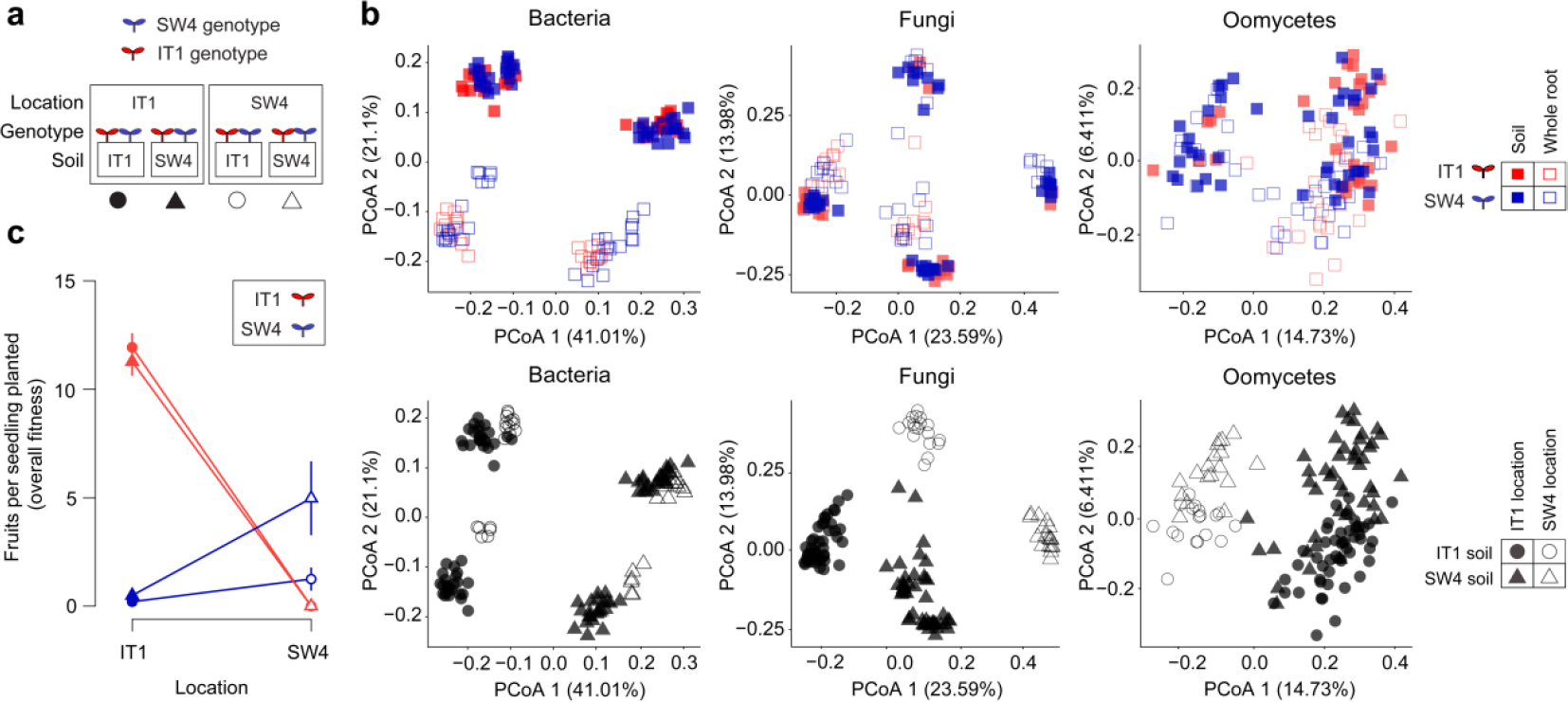
Reciprocal transplant between two *A. thaliana* populations in Sweden and Italy. **a**, Schematic overview of the reciprocal transplant experiment. Soils and plant genotypes from IT1 and SW4 sites were reciprocally transplanted in the two locations (eight different treatment combinations). The symbols below the schematic view correspond to the symbols also used in the other panels. **b**, Community structures of bacteria, fungi, and oomycetes in the 131 samples were determined using principal coordinate analysis (PCoA). The first two dimensions of the PCoA are plotted based on Bray-Curtisdistances. To facilitate visualization, the same PCoA plot was represented either according to the genotype and the compartment (upper panels) or according to the soil origin and the location (lower panels). Note that no Italian plant survived at the Swedish site. **c**, Overall fitness (number of fruits per seedling planted; mean ± SE) of Italian and Swedish genotypes (red and blue color, respectively) when reciprocally planted in Italian and Swedish soils (circle and triangle symbols, respectively) and grown at Italian and Swedish locations (filled and open symbols, respectively).

Consistent with earlier reports, a significant proportion of the variance remained unexplained in bacterial, fungal, and oomycetal communities in both soil and root compartments (> 40%, data not shown), likely arising from unmeasured environmental variables, stochastic processes, or species interactions such as microbe-microbe interactions. To identify signatures of microbial interactions, we quantified correlations between bacterial and fungal alpha diversity across sites. The correlation between bacterial and fungal diversity was positive in soil samples (Spearman’s rank correlation, observed OTUs: r_s_ = 0.30, p < 0.001, Shannon index: r_s_ = 0.21, p = 0.002), but negative in root endosphere samples (Observed OTUs: r_s_ = −0.15, p = 0.092, Shannon index: r_s_ = −0.24, p = 0.005) (**Supplementary Fig. 6a, b**). The observation that bacterial diversity is negatively correlated with fungal diversity in the root endosphere suggests that microbial interactions also contribute to community differentiation at the soil-root interface, as recently reported^3^. Overall, our results suggest that among-site variation in environmental conditions affected root-associated microbial communities more strongly than did temporal variation within sites. In contrast, differences between host species had less impact on root microbiota differentiation than had spatial and temporal variations. Therefore, differences in soil properties (e.g., pH, which ranged from 5.1 to 7.9) and climate are likely causes of the variation in root microbiota among European populations of *A. thaliana*.

### Site-specific differences in soil, climate, and genotype drive root microbiota differentiation between two *A. thaliana* populations

We hypothesized that continental-scale variation in the *A. thaliana* root microbiota is mediated by interactions between soil, climate, and host genotype and that these factors might differentially influence the establishment of bacteria and filamentous eukaryotes in plant roots. To disentangle the respective contribution of these three factors on microbial community composition at the root interface, we conducted a transplant experiment between two geographically widely separated *A. thaliana* populations in Sweden and Italy (SW4 and IT1) (**Supplementary Fig. 7a, b**). In this experiment, we transplanted seedlings of each plant genotype into soils from both SW4 and IT1 at each site in autumn, at the time of natural seedling establishment in the local populations (**Fig. 4a**). At fruit maturation, the two *A. thaliana* genotypes were harvested and community composition was defined for bacteria, fungi and oomycetes in soil (Soil with RS-associated microbes) and whole root (Root-with RP-associated microbes) samples (n = 131, **Fig. 4b**).

Consistent with the European transect experiment, we observed a greater compartment effect for bacteria than for filamentous eukaryotes, explaining 17.8%, 7.4%, and 3.1% of the variance in bacterial, fungal, and oomycetal community composition, respectively (PERMANOVA, p < 0.001, **Fig. 4b** and **Supplementary Table 7**). However, the degree of structural convergence in roots was weaker, probably due to the fact that we did not partition the RP compartment from the root endosphere in this experiment. Because no Italian plants survived at the Swedish site, we used a canonical analysis of principal coordinates (CAP; capscale function) to assess the effect of genotype among whole root samples at the Italian site only. CAP analysis constrained by genotype indicated a significant effect of host genotype for all three microbial groups in plant roots, explaining 2.4% (p = 0.001), 2.36% (p = 0.026) and 2.9% (p = 0.002) of the variance in bacterial, fungal, and oomycetal community composition, respectively (**Supplementary Fig. 7d**).

The origin of soil (Italy *vs.* Sweden) explained substantially more variation in bacterial soil biota than did transplant location (soil origin: 62.3%, transplant location: 15.8%; PERMANOVA, p < 0.001). This was also the case for bacterial communities associated with whole root samples (soil origin: 47.1%, transplant location: 17.4%; PERMANOVA, p < 0.05) (**Fig. 4b** lower panel, **Supplementary Table 7)**. The percentage of variation explained by the origin of soil was weaker for fungi and oomycetes in the soil fraction (fungi: 29.6%, oomycetes: 18.54%; PERMANOVA, p < 0.001), whereas differences other than soil origin, including climatic differences between sites, also largely accounted for between-site variation in microbial eukaryotic assemblages in soil (fungi: 27.5%, oomycetes: 6.3%; PERMANOVA, p < 0.001). In whole root samples, the effect of the location was as strong as the effect of the soil for fungi (location: 14.6%, soil origin: 15.1%; PERMANOVA, p < 0.001) and stronger than the effect of the soil for oomycetes (location: 10.6%, soil origin: 4.7%; PERMANOVA, p < 0.001) (**Fig 4b** lower panel, **Supplementary Table 7**). These results indicated that soil origin and transplant location differentially affect the assembly of root-associated bacteria and filamentous eukaryotes.

Inspection of root-associated microbial OTUs that responded to differences in soil origin and/or location in the reciprocal transplant demonstrated the stable prevalence of geographically widespread bacterial OTUs in root samples representing different soil × location × genotype combinations (**Supplementary Fig. 8a, b** and **c)**. Abundance profiles of microbial taxa in whole root samples were impacted by soil origin, location, and their interaction, with notable differences depending on microbial classes. For example, variation in root-associated Actinobacteria was almost exclusively explained by soil origin, whereas variation in root-associated Alpha-, Beta-, and Gamma-proteobacteria was explained by soil origin and location to similar degrees (**Supplementary Fig. 8d**). Similarly, the presence of root-associated fungi belonging to Dothideomycetes was primarily explained by location, whereas the presence of root-associated fungi belonging to Leotiomycetes was largely explained by soil origin (**Supplementary Fig. 8d**). The results point to differential impact of site and soil specific factors on different taxonomic groups of the root microbiota. Overall, 74.4% and 86.3% of root-associated fungal and oomycetal OTUs, respectively, responded to difference in location or location x soil, whereas this percentage was reduced to 44.7% for root-associated bacterial OTUs (**Supplementary Fig. 8d)**. Taken together, our data suggest that location-specific factors such as climatic conditions affect differentiation of root-associated filamentous eukaryotic communities more than that of bacteria, and this likely explains variation in community structure observed for these microbes among European sites and across successive years (**Fig. 3** and **Supplementary Fig. 4b**).

### Site-specific differences in soil conditions only weakly contributes to adaptive differentiation between *A. thaliana* populations

If adaptation to the biotic or abiotic characteristics of the local soil contributes to adaptive differentiation between *A. thaliana* populations, the advantage of the local over nonlocal genotype should be greater when plants are grown on local soil than when grown on non-local soils. To test this hypothesis, we scored plant survival and fecundity (number of fruits per reproducing plant) and estimated overall fitness (number of fruits per seedling planted) of the plants grown in the reciprocal transplant experiments at the sites of IT1 and SW4 (n = 1,008; **Supplementary Table 8**).

At the IT1 site the relative fitness of the local genotype was higher on local than on non-local soil: the mean fitness of the Italian genotype was 58 times higher than that of the non-local Swedish genotype on Italian soil (corresponding to a selection coefficient of s = 0.98), and 23 times higher (s = 0.96) on Swedish soil (significant soil x genotype interaction, F_1,10_ = 5.7, p = 0.038; **Fig. 4c, Supplementary Table 9**). The strong local advantage was a function of both higher survival and fecundity (**Supplementary Fig. 7c**). At the SW4 site, no Italian plant survived to reproduce, meaning that selection coefficients were 1 on both soil types (**Fig. 4c**). It was therefore not meaningful to compare adaptive differentiation on the two soil types. However, the Swedish genotype showed 3.5-fold higher survival and 4.0-fold higher overall fitness when planted in Swedish soil compared to in Italian soil (survival: one-tailed test, t = 1.81, p = 0.060; overall fitness: t = 2.14, p = 0.039) **(Fig. 4c, Supplementary Fig. 7c)**. These results demonstrate a strong selective advantage to the local *A. thaliana* genotype at both sites, consistent with previous studies on the same populations^15, 28^. However, despite the marked differences in the geochemical properties and microbial communities of the soil, adaptation to local soil conditions explained only a small fraction of the adaptive differences between the two *A. thaliana* populations. This suggests that adaptation to climatic variables not related to the characteristics of the soil are the primary drivers of adaptive differentiation between *A. thaliana* populations in northern and southern Europe.

## Discussion

By monitoring the root microbiota of *A. thaliana* in 17 natural populations across three successive years, we observed strong geographic structuring of soil microbial communities, but a surprising degree of convergence among root-associated taxa. The convergence in microbial community composition observed for root endosphere samples is remarkable given the large geographical distances between sites, contrasting edaphic characteristics, and distinct microbial communities in the surrounding bulk soils. Our results indicate that environmental factors and variables that explain geographic distribution of microbes in soil have less predictive power in plant roots. The degree of convergence in community structure varied among microbial groups and was more evident for the bacterial root microbiota than for filamentous eukaryotes. Differences between sites that were independent of soil characteristics largely explained differentiation in filamentous eukaryotic communities in our reciprocal transplant experiment, suggesting a link between climatic conditions and variation observed for filamentous eukaryotic communities across sites and years. Our results indicate that difference in climate and soil between sites not only contribute to variation in root-associated microbial communities, but also to adaptive divergence between two *A. thaliana* populations in northern and southern Europe.

The striking structural convergence observed for bacterial communities in roots of *A. thaliana* and grasses was explained by the presence of a few, diverse and geographically widespread taxa that disproportionately colonize roots across sites. The conserved enrichment of Beta- and Gamma-proteobacteria in *A. thaliana* roots, together with the identification of 13 habitat-generalist OTUs that were consistently and abundantly detected in the roots of *A. thaliana* and grass populations suggest potential co-evolutionary histories between these microbes and evolutionary distant plant species. These few OTUs belong to bacterial genera that are often detected in plant roots across various environmental conditions^29, 30, 31, 32, 33^. Notably, *Bradhyrhizobium* and *Burkholderia* (OTUs 8, 4987, 13, 14, and 4486 in this study) were the two most dominant of 47 widespread genera in roots of phylogenetically diverse flora across a 10 km transect in Australia^32^. The low number of widespread core bacterial taxa detected in our study might be due to the large distances between sites and extensive differences in soil pH (ranging from 5.1 to 7.9), contrasting with the more uniform range of soil pH reported in ref.^32^ (i.e. 4.1 to 4.6). Our results nonetheless suggest that at least part of the similarity in bacterial community composition observed in roots of divergent plant species in microbiome studies^**34**^ is driven by the presence of geographically widespread taxa that efficiently colonize plant roots across a broad range of environmental conditions.

The remarkable phylogenetic diversity among the few geographically widespread bacteria detected in plant roots suggests convergent evolution and metabolic adaptation to the root habitat in phylogenetically distant bacterial lineages^35^. Similar to what has been described in leaves for wild *A. thaliana*^36^, a Pseudomonas OTU in our study was also by far the most dominant taxon in *A. thaliana* roots (i.e. OTU5, Relative abundance in roots = 14.4% in average), pointing to Pseudomonas taxa as the most successful and robust colonizers of wild *A. thaliana*. Although we sampled healthy-looking plants in their natural habitats, we cannot exclude the possibility that some of these bacterial strains represent widespread pathogens that negatively affect plant fitness without causing disease symptoms in nature^36^. Alternatively, these taxa might carry out widespread and important beneficial functions for *A. thaliana* survival such as pathogen protection e.g., *Pseudomonas, Pelomonas, Acidovorax, Flavobacterium*^3, 35, 37^, abiotic stress tolerance, or plant growth promotion (e.g., *Bradyrhizobium, Burkholderia*^38, 39^). It remains to be seen whether this widespread association between plant roots and bacteria is evolutionary ancient and the extent to which it has contributed to plant adaptation to terrestrial ecosystems. Irrespective of this, the 13 core OTUs identified here in *A. thaliana* roots in natural environments provide a rational framework for future design of low complexity synthetic bacterial communities from culture collections of root commensals and co-culturing with gnotobiotic plants^40^ to study their contribution to plant health and growth in laboratory environments.

The results of the reciprocal transplant between two widely separated *A. thaliana* populations (one in Italy and one in Sweden; IT1 and SW4) demonstrated significant effects of soil and location on microbiota assembly in both soil and root compartments. Our observation that transplant location affected the community composition of filamentous eukaryotes more strongly than that of bacteria suggest that the response to climate varies among microbial kingdoms^41, 42^. Our data corroborate the hypothesis that climate is a key driver of among-site variation and geographic distribution of filamentous eukaryotes in soil, as predicted based on association studies^13, 43, 44^. This contrasts with the geographic distribution of soil bacteria which is known to be primarily controlled by edaphic factors^10^. The marked among-year variation in community composition of filamentous eukaryotes shows that year-to-year differences in environmental conditions can significantly modify geographical patterns of filamentous microbes at the soil-root interface across Europe. Our field transplant experiment also demonstrated an effect of host genotype (Italian vs. Swedish accession) on the composition of root microbial communities. The proportion of variation in community composition explained by host genotype was limited (∼ 2% at OTU resolution) compared to that explained by origin of soil and location of experiment, but is still comparable to estimates for host genotype obtained in previous studies of *A. thaliana* (ref.^45^; variance: < 1%, p < 0.05), *Boechera stricta* (ref.^46^; genotype effect not significant), and *Populus* (ref.^47^; variance: ∼ 3%, p < 0.05). Taken together, our results suggest that bacterial and fungal assemblages in roots are differentially controlled by edaphic and climatic conditions, and that host genetic differences contribute only little to root microbiota differentiation between the two *A. thaliana* populations.

Local adaptation is common among plant populations, but the extent to which divergent selection and local adaptation can be attributed to local soil conditions has been examined mainly in relation to high concentrations of heavy metals^48^, serpentine^49^, and high salinity^25, 50^. The field experiment of the present study, in which both soils and plant genotypes were reciprocally transplanted at two locations in Sweden and Italy, demonstrated strong local adaptation between two geographically widely separated *A. thaliana* populations. The relative performance of local and non-local host genotypes was primarily affected by the geographical location and only weakly by soil origin, despite extensive differences in microbial community composition and physical and chemical properties between Swedish and Italian soils (**Fig. 1d** and **Supplementary Table 1**). Particularly, differences in available Ca, reserve K, available Mg, pH, iron, and available K between the two soils accounted for 99.4%, 93.3%, 87.9%, 60.7, 39.4% and 30%, of the total variation, respectively, observed for these parameters across all 17 soils (**Supplementary Table 1**). In their native habitats, the two populations are exposed to widely different climates and also to differences in seasonal changes of day length. Previous work has demonstrated that the relative survival of the Italian genotype in Sweden is negatively related to minimum soil temperature in winter^15^, and also that genetic differences in phenological traits such as timing of germination and flowering can explain a substantial portion of selection against the non-local genotype at the two sites^**51, 52, 53**^. Taken together, our data indicate that differences in climate have been more important than differences in soil and endogenous microbiota for the adaptive divergence between the two study populations. Although we did not uncouple soil-mediated from microbe-mediated local adaptation in this study, our results suggest that the extensive differences in microbial community composition between IT1 and SW4 soil contribute little to adaptive differentiation between two *A. thaliana* populations in northern and southern Europe. Future studies should determine whether the environmental factors that affect root microbiota assembly and adaptive differentiation at large spatial scales act differentially at smaller geographical scales.

## Methods

### Harvesting of *A. thaliana* and grasses across 17 European sites

We selected seventeen sites with variable soil characteristics across a climatic gradient from Sweden to Spain. At each site, *A. thaliana* and sympatric grasses occur naturally^54, 55, 3, 1, 15^ (**Supplementary Table 1, Fig. 1a**). We harvested *A. thaliana* from February to May at the same developmental stage (bolting/flowering stage), for one (three sites), two (one site) and three (thirteen sites) consecutive years (**Supplementary Table 1**). Plants were harvested with their surrounding bulk soil with a hand shovel without disturbing the plant root system, transferred to 7×7 cm greenhouse pots and transported immediately to nearby laboratories for further processing. Four plant individuals pooled together were considered as one pooled-plant technical replicate (4 technical replicates in total). In addition, four plants were not pooled and kept individually as single-plant replicates to assess difference in microbial community composition between single plant and pooled plants. Note that these two sampling strategies explain only 0.8, 3 and 2.4% of the variation in bacterial, fungal, and oomycetal community composition, respectively (**Supplementary Table 5, Supplementary Fig. 4b**). At each of the 17 sites, we also harvested and processed similarly three neighbouring plants growing within 50 cm of *A. thaliana* and belonging to the Poaceae family. In total, we collected 285 plants across three years.

### Fractionation of soil and root samples

To distinguish and separate four microbial niches across the soil-root continuum, plants and respective roots were taken out from the pot. Samples from the bulk soil exempt from root or plant debris were taken, snap-frozen in liquid nitrogen and stored for further processing (Soil compartment). Individual plants were manually separated from the main soil body and non-tightly adhered soil particles were removed by gently shaking the roots. These roots were separated from the shoot and placed in a 15-mL falcon containing 10 mL of deionized sterile water. After 10 inversions, roots were transferred to another falcon and further processed, while leftover wash-off (containing the RS fraction) was centrifuged at 4,000 g for 10 min. Supernatant was discarded and the pellet was resuspended and transfered to a new 2-mL screw-cap tube. This tube was centrifuged at 20,000 rpm for 10 minutes, the supernatant was discarded and the pellet was snap-frozen in liquid nitrogen and stored for further processing (RS compartment). After RS removal, cleaned roots were placed in a 15-mL falcon with 6 mL of detergent (1xTE + 0.1% Triton® X-100) and manually shaken for 2 minutes. This step was repeated for a total of three detergent washes, in between which, roots were transferred to a new 15-mL falcon with new detergent. After these washes in detergent, roots were transferred to a new 15-mL falcon. The remaining washes (approximately 18 mL) were transferred to a 20 mL syringe and filtered through a 0.22 µM-pore membrane. The membrane was snap-frozen in liquid nitrogen until further processing (RP compartment). Lastly, three-times washed roots were subjected to a further surface sterilization step to further deplete leftover microbes from the root surface. Roots were sterilized one minute in 70% ethanol, followed by one minute in 3% NaClO, and rinsed five times with deionized sterile water. These roots were dried using sterile Whatman paper and snap-frozen in liquid nitrogen until further processing (Root compartment) (**Supplementary Fig. 1a**).

In total, 1,125 samples were produced after fractionation. We validated the fractionation protocol by printing processed roots on Tryptic Soy Agar (TSA) 50% before fractionation, after each detergent wash and after surface sterilization. Washes produced after each fractionation step (without treatment, after each detergent step and after surface sterilization) were also plated on TSA 50%. In brief, treated roots were cut and placed on plates containing TSA 50% medium for 30 sec and then removed. After 3 days at 25 °C, colony forming units (CFU) were counted. Similarly, 20 µL of remaining washes were spread onto TSA 50% medium-containing plates and CFU were counted after 3 days of incubation at 25 °C (**Supplementary Fig. 1b**).

### Reciprocal transplant experiment

We reciprocally transplanted both soil and local *A. thaliana* genotypes between sites in Italy (population IT4) and Sweden (population SW4; **Supplementary Fig. 7a).** See ref.^15^ for a description of study sites and plant genotypes. Soil was collected at the two experimental sites in spring 2016 and stored at 6°C until the establishment of the experiments. Seeds were planted in Petri dishes on agar, cold stratified in the dark at 6°C for one week, and then moved to a growth room (22°C/16°C, 16 h day at 150 μE/m2/sec PAR, 8 h dark) where the seeds germinated. Nine days after germination, seedlings were transplanted to 299-cell plug trays (cell size: 20 mm × 20 mm × 40 mm) with local and non-local soils in blocks of 6 × 7 cells in a checkerboard design (three blocks of each soil type per tray). At each site, we transplanted 20 local and 22 non-local seedlings into randomized positions in each of six blocks per soil type, with a total of 120 local and 132 non-local plants for each site × soil combination. During transplantation, plug trays were kept in a greenhouse at about 18°C/12°C and 16 h day/8 h night. Within six days, the trays were transported to the field sites where they were sunk into the ground (on 9 September 2016 in Sweden, and on 29 October 2016 in Italy). The transplanted seedlings were at approximately the same stage of development as naturally germinating plants in the source population.

At fruit maturation in spring 2017, we scored survival to reproduction and number of fruits per reproducing plant (an estimate of fecundity), and estimated total fitness as the number of fruits produced per seedling planted (following ref.^15^). Statistical analyses were based on block means for each genotype. No Italian plant survived at the Swedish site, and it was therefore not possible to fit a model examining the site × soil × genotype interaction. Instead, we conducted analyses separately by site. For the Italian site, we assessed differences in overall fitness due to soil, genotype, and the soil × genotype using a mixed-model ANOVA, with block (random factor) nested within soil type. In Sweden, we used a one-tailed t-test to test the prediction that survival and overall fitness of the Swedish genotype is higher when planted in Swedish than when planted in Italian soil.

At the time of fitness assessment, we harvested plants and their surrounding soil by removing the whole soil plug. Soil was separated from the roots manually and a soil sample was taken. After removing loose soil particles by gentle shaking, roots were placed in a 15-mL falcon and washed by inverting with 10 mL of deionized water and surface-sterilized by washing for one minute with 70% ethanol, then washing with 3% NaClO for one minute and rinsed several times with deionized sterile water. These whole root samples correspond to the combination of Root and RP compartments described for the European transect survey. Six to twelve soil samples were harvested for each of the eight combinations of soil, location and genotype (n = 72), as well as three to twelve whole root samples per condition (n = 59). Note that no Italian plants survived at the Swedish site. In total, 131 root and soil samples were harvested, stored in dry ice, and processed for DNA isolation and microbial community profiling.

### Microbial community profiling

Total DNA was extracted from the aforementioned samples using the FastDNA SPIN Kit for Soil (MP Biomedicals, Solon, USA). Samples were homogenized in Lysis Matrix E tubes using the Precellys 24 tissue lyzer (Bertin Technologies, Montigny-le-Bretonneux, France) at 6,200 rpm for 30 s. DNA samples were eluted in 60 μL nuclease-free water and used for bacterial, fungal and oomycetal community profiling^1, 3^. The concentration of DNA samples was fluorescently quantified, diluted to 3.5 ng/µL, and used in a two-step PCR amplification protocol. In the first step, V5–V7 of bacterial 16S rRNA (799F - 1192R), V2-V4 of bacterial 16S rRNA (341F – 806R), fungal ITS1 (ITS1F - ITS2), fungal ITS2 (fITS7 – ITS4) and oomycetal ITS1 (ITS1-O - 5.8 s-Rev-O) (**Supplementary Table 2**) were amplified. Under a sterile hood, each sample was amplified in triplicate in a 25 μl reaction volume containing 2 U DFS-Taq DNA polymerase, 1x incomplete buffer (both Bioron GmbH, Ludwigshafen, Germany), 2 mM MgCl_2_, 0.3% BSA, 0.2 mM dNTPs (Life technologies GmbH, Darmstadt, Germany) and 0.3 μM forward and reverse primers. PCR was performed using the same parameters for all primer pairs (94°C/2 min, 94°C/30 s, 55°C/30 s, 72°C/30 s, 72°C/10 min for 25 cycles). Afterward, single-stranded DNA and proteins were digested by adding 1 μl of Antarctic phosphatase, 1 μl Exonuclease I and 2.44 μl Antarctic Phosphatase buffer (New England BioLabs GmbH, Frankfurt, Germany) to 20 μl of the pooled PCR product. Samples were incubated at 37°C for 30 min and enzymes were deactivated at 85°C for 15 min. Samples were centrifuged for 10 min at 4000 rpm and 3 μl of this reaction were used for a second PCR, prepared in the same way as described above using the same protocol but with cycles reduced to 10 and with primers including barcodes and Illumina adaptors (**Supplementary Table 2**). PCR quality was controlled by loading 5 μl of each reaction on a 1% agarose gel. Afterwards, the replicated reactions were combined and purified: 1) bacterial amplicons were loaded on a 1.5% agarose gel and run for 2 hours at 80 V; bands with the correct size of ∼500 bp were cut out and purified using the QIAquick gel extraction kit (QIAGEN, Hilden, Germany); 2) fungal and oomycetal amplicons were purified using Agencourt AMPure XP beads. DNA concentration was again fluorescently determined, and 30 ng DNA of each of the barcoded amplicons were pooled in one library per microbial group. Each library was then purified and re-concentrated twice with Agencourt AMPure XP beads, and 100 ng of each library were pooled together. Paired-end Illumina sequencing was performed in-house using the MiSeq sequencer and custom sequencing primers (**Supplementary Table 2**).

### 16S rRNA gene and ITS read processing

All paired rRNA amplicon sequencing reads have been analysed with a pipeline described recently^3^. The main parts include scripts from QIIME^56^ and usearch^57^. OTUs were clustered at a 97% threshold (usearch – cluster otus). Bacterial reads were checked against the greengenes database^58^ to remove non-bacterial reads. Fungal and oomycetal reads were checked against an ITS sequences database (full length ITS sequences from NCBI) to remove unwanted reads. Taxonomic assignment was done via QIIME (assign_taxonomy) for bacterial OTUs, using the greengenes database. Taxonomic assignment for fungal OTUs was done via RDP-classifier^**59**^ against the Warcup database^60^. Assignment of oomycetal OTUs was also done via RDP but against a self-established database. For rRDP-based classification a cut-off of 0.5 was used to filter out uncertain assignments.

### Computational analyses

All OTU tables obtained from the before mentioned pipelines were filtered for very low abundant OTUs prior any analysis and post processing. For this, only OTUs having at least 0.1% RA in at least one sample were kept. These filtered tables were used for all further analyses. For OTU based analysis of bacterial data, OTUs assigned as chloroplast- or mitochondria-derived were removed prior to analysis. Alpha diversity indices (Shannon index and observed OTUs) were calculated using OTU tables rarefied to 1,000 reads. Bray Curtis distances between samples were calculated by using OTU tables that were normalized using cumulative-sum scaling (CSS^**61**^). Average Bray-Curtis distances across replicates and years, were calculated by using the mean across all combinations between two sets of sites (e.g. between all soil samples from site x and site y). Comparisons of samples coming from different years were not considered for this analysis. These average distances were hierarchically clustered (hclust in R, method = “complete”). Similarly, averaged relative abundances (avg. RA)of taxonomic groups was achieved by averaging across all samples from a particular site – compartment combination. All principal coordinate analyses were calculated using the respective Bray-Curtis distance matrices as an input for cmdscale function (standard R). Explained variance of different factors was based on Bray-Curtis distances. These were used as an input for PERMANOVA analysis using the adonis function in R (vegan package). All factors have been analysed independently. Dependency of data was either tested via a linear regression (using lm function from R) or Spearman rank correlation (cor.test function in R). Unless noted differently, a p value < 0.01 was considered significant. Differential abundance of taxonomic groups was tested using the Wilcoxon rank sum test (wlicox.test in R, FDR < 0.05).

For calculating site prevalence of OTUs, OTU tables were restricted to samples having > 1,000 reads. Count tables were transformed to relative abundances by division of total column (sample) sums. An OTU was marked as present in a given sample if its RA was > 0.1%. OTUs present in < 5 samples were not considered further. Site prevalence reflects in how many sites (out of all European sites sampled in a given year) an OTU was present on average across years (here only years where an OTU was detected were taken into account). Site prevalence for OTUs present in neighbouring plants was calculated in the same way. To calculate the average RA across sites and years for a given OTU, only those samples were the OTU was actually present were considered. In this way, prevalence and mean RA are independent. OTUs present in > 80% of sites on average were considered to be geographically widespread. Differential abundance of OTUs in the different groups (geographically restricted, common, and widespread) was calculated between compartment using Kruskal-Wallis test (kruskal.test with dunn.test in R, FDR < 0.05). Root-specific enrichment for widespread OTUs, compared to soil and RS samples was tested with a generalized linear model, as described in ref.^22^ (FDR, p < 0.05). To compare generalist OTUs from this study with OTUs found in roots of *Lotus japonicus*^26^ bacterial OTUs were directly compared using the representative sequences (Blastn, 98% sequence identity). To compare fungal OTUs, the representative sequences from geographically widespread OTUs were blasted against full ITS sequences from the UNITE database^62^. The best hits received were then used to find similar OTUs in the *L. japonicus* data (98% sequence identity).

Enrichment patterns of root associated OTUs in the transplant experiment (see heatmap in **Supplementary Fig. 8**) were examined as follows. For bacterial and fungal datasets, samples with less than 1,000 reads were removed, whereas for the oomycetal dataset, samples with less than 250 reads were removed. OTUs having a mean relative abundance > 0.01% across all root samples were kept for the analysis. Relative abundance entries of zero were replaced by 0.001%. All relative abundance values were log2 transformed and these data were used as input for generating a heatmap (heatmap.2 function in R, gplots library). Enrichment in one of the six tested conditions (Swedish location, Italian location, Swedish soil, Italian soil, Italian location and soil, Swedish location and soil) was estimated by comparing the mean relative abundance of each OTU across all samples with the mean relative abundance in respective sample combinations. E.g. to be enriched at the Italian location, an OTU must be more abundant at the Italian location compared to the Swedish location, irrespective of the soil (see panel in **Supplementary Fig. 8)**. OTUs that are not enriched in any of the conditions but are prevalent across samples (mean relative abundance above 0.1%) are marked in grey in the heatmap. OTUs that were not prevalent and not enriched in any condition are not shown. Geographically widespread OTUs were identified by comparing representative sequences from the transect data with those of the reciprocal transplant experiment (Blast at 98% sequence identity).

## Data availability

Sequencing reads of samples from the European transect experiment and the reciprocal transplant experiment (MiSeq 16S rRNA and ITS reads) have been deposited in the European Nucleotide Archive (ENA) under accession numbers ENA: ERP115101 and ENA: ERP115102, respectively.

## Code availability

All scripts for computational analysis and corresponding raw data are available at https://github.com/ththi/European-Root-Suppl.

### Acknowledgements

We thank Neysan Donnelly for scientific English editing. This work was supported by funds from the Max Planck Society to S.H. and P.S-L., a European Research Council starting grant (<u>MICRORULES</u>) to S.H., a European Research Council advanced grant (<u>ROOTMICROBIOTA</u>), and the “Cluster of Excellence on Plant Sciences” program funded by the Deutsche Forschungsgemeinschaft to P.S.-L., and grants from the Swedish Research Council to JÅ. C.A.-B. lab was funded by grant BIO2016-75754-P (AEI/FEDER).

## Author contributions

S.H., P.S-L. and J.Å. conceived the project. E.K., F.R., C.A-B., J.Å., and S.H. identified natural. *A. thaliana* populations. P.D. and S.H. collected the samples. P.D. prepared all samples and performed microbial community profiling. P.D. and T.T analyzed microbiota data. T.E, and J.Å. prepared the field reciprocal transplant experiment. J.Å., T.E., and P.D. analyzed plant fitness data. S.H. supervised the project. T.T., P.D., J.Å., P.S-L., and S.H. wrote the paper.

## Supplementary Figures

**Supplementary Fig. 1:**
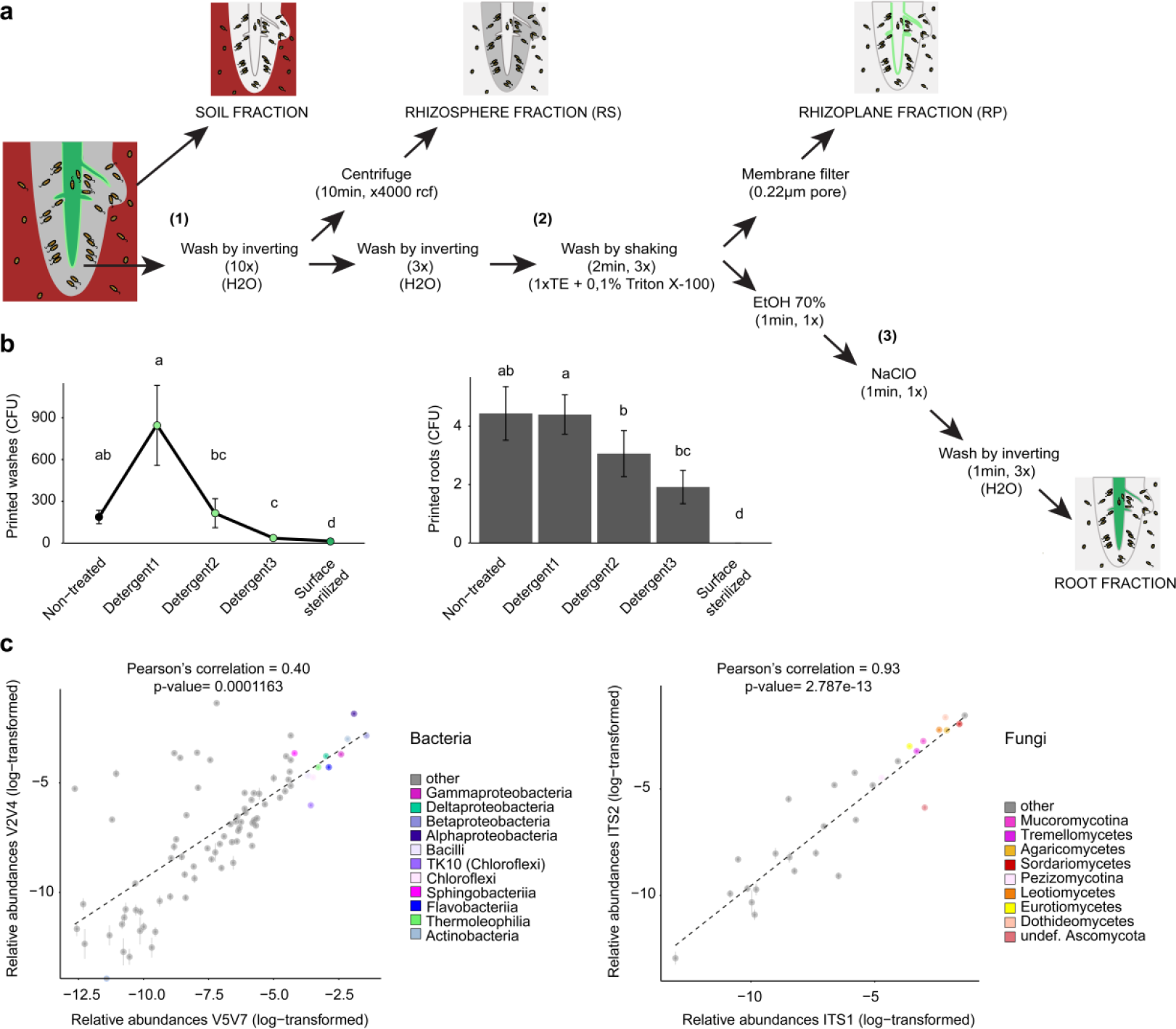
Validation of the root fractionation protocol and assessment of primer amplification bias. **a**, Protocol to fractionate four microbial niches across a distance gradient from bulk soil to roots’ interior. Roots of *A. thaliana* grown in their natural environments were briefly washed (1) to separate loosely attached soil particles from the root surface (rhizosphere, RS). After a second washing step, roots were vigorously washed with detergent (three times) to capture microbes that tightly adhere to the root surface (2). The resulting washes were then filtered through a 0.22 µM membrane (rhizoplane, RP). Finally, surface sterilization of root samples by consecutive EtOH and NaClO washes enriched the final root sample in microbial root endophytes (3). **b**, Validation of the fractionation protocol (depicted in panel a) was performed by printing root washes (left panel) and washed roots (right panel) on 50% Tryptic Soy Agar medium. Sequential detergent washes efficiently release microbes from the root surface and further root surface sterilization prevents the growth of rhizoplane-associated microbes (Wilcoxon rank sum test, p < 0.01). All three detergent steps were combined and filtered to prepare the RP fraction (light green dots, left panel). **c**, Comparison of bacterial (left panel) and fungal (right panel) classes profiled with the V2V4 and V5V7 regions of the bacterial 16s rRNA gene, and the ITS1 and ITS2 of the fungal ITS. The correlation between the relative abundances of each microbial class is shown (Pearson’s correlation, p < 0.001).

**Supplementary Fig. 2:**
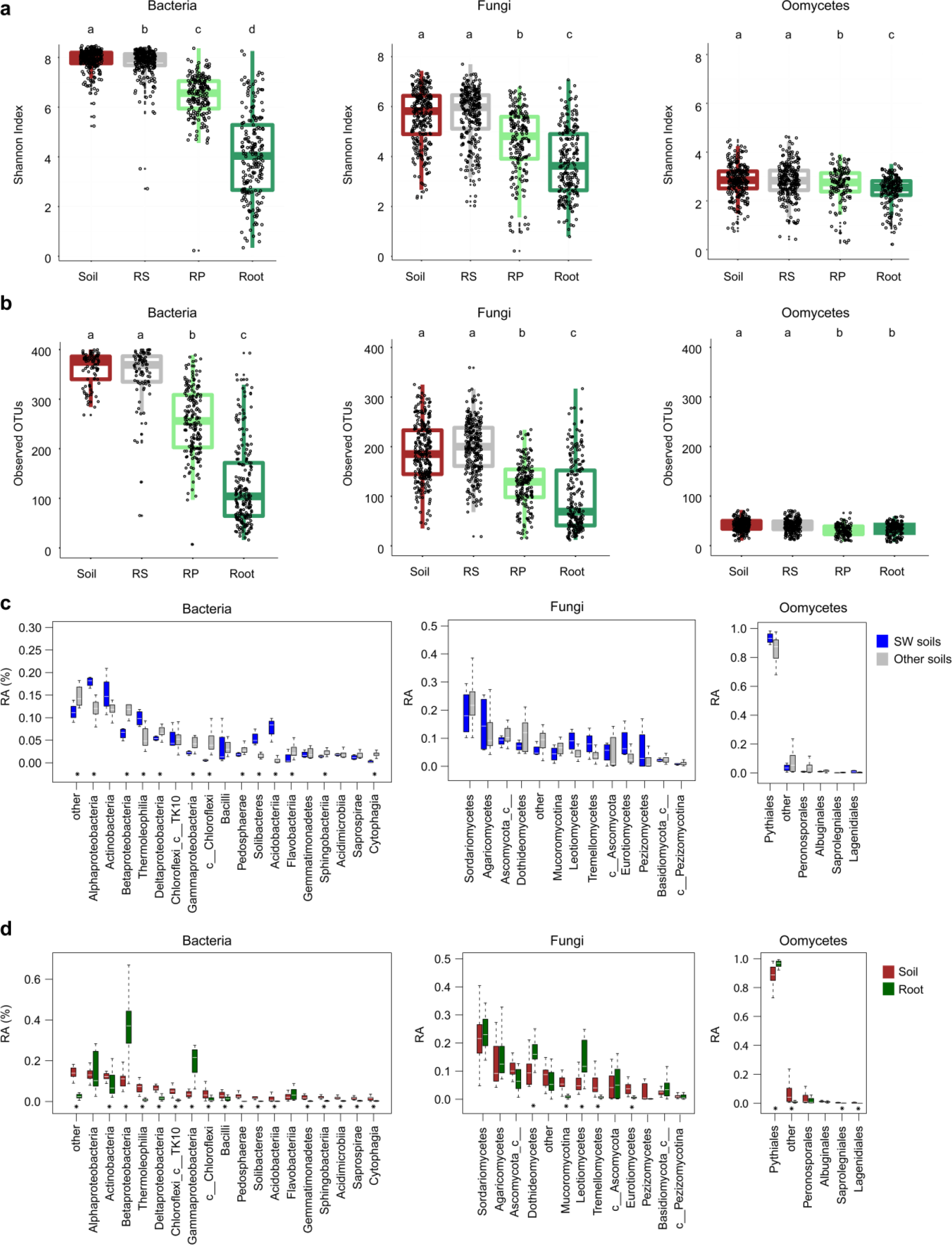
Microbial alpha diversity and enrichment signatures in plant roots. **a**, Microbial alpha diversity measured across all 17 sites in soil, rhizosphere (RS), rhizoplane (RP), and root samples based on the Shannon index. All samples from a given site were taken into account and the datasets were rarefied to 1,000 reads. Individual data points within each box correspond to samples from the 17 natural sites (Kruskal-Wallis test, p < 0.05). **b**, Microbial alpha diversity measured across all 17 sites in soil, rhizosphere (RS), rhizoplane (RP), and root samples based on the number of observed OTUs. All samples from a given site were taken into account and the datasets were rarefied to 1,000 reads. Individual data points within each box correspond to samples from the 17 natural sites (Kruskal-Wallis test, p < 0.05). **c**, Comparison of taxa relative abundance (RA) between Swedish soils (SW1-4, blue) and the other European soils (grey) for bacteria (left), fungi (middle), and oomycetes (right). RA is aggregated at the class (bacteria and fungi) and order (oomycetes) levels and significant differences are marked with an asterisk (Wilcoxon rank sum test, FDR < 0.05). **d**, Comparison of taxa RA between soil (dark red) and root (dark green) samples for bacteria (left), fungi (middle), and oomycetes (right). RA measured in soil and root samples across the 17 *A. thaliana* populations were aggregated at the class (bacteria and fungi) and order (oomycetes) levels. Significant differences are marked with an asterisk (Wilcoxon rank sum test, FDR < 0.05).

**Supplementary Fig. 3:**
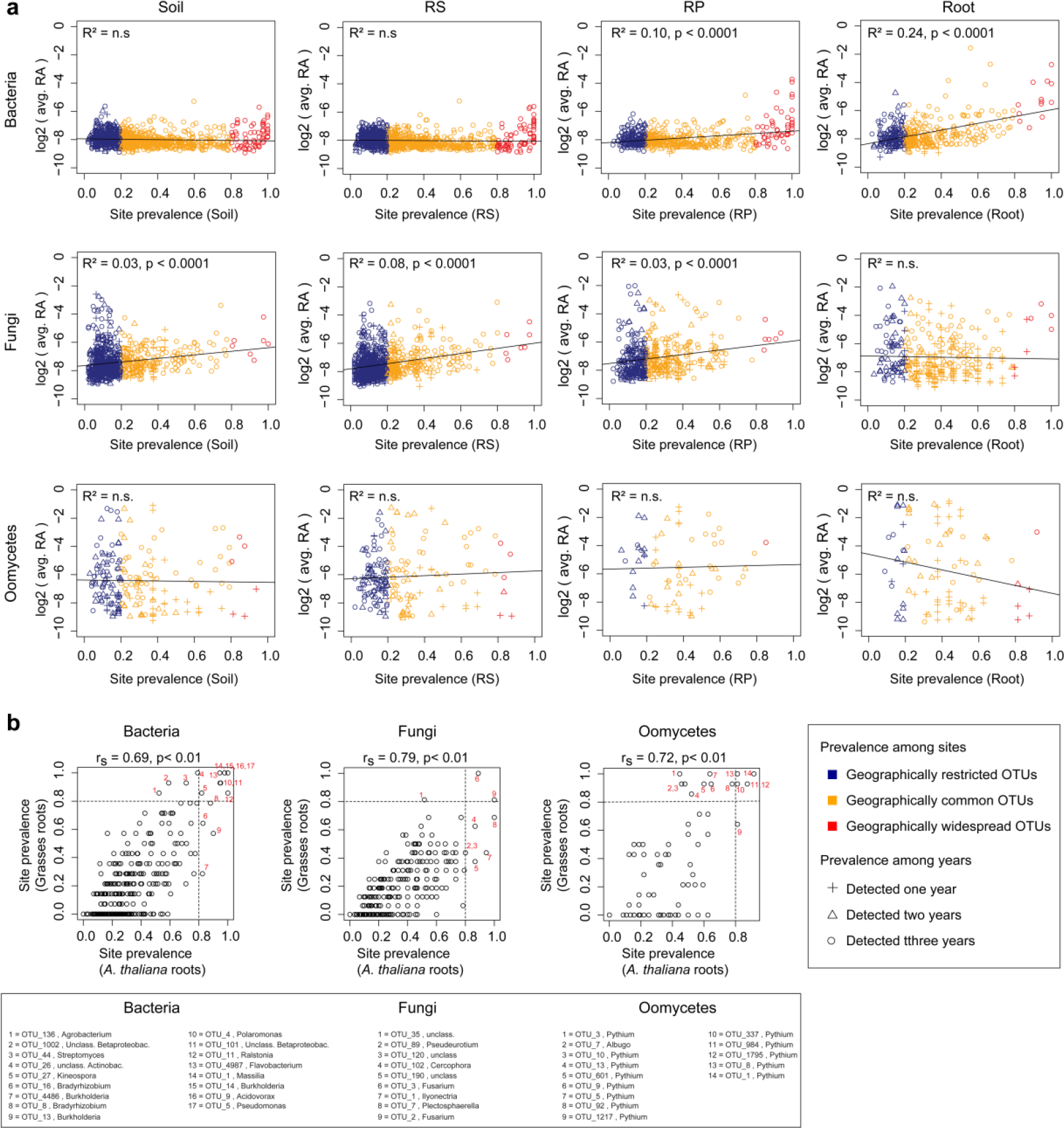
Geographically widespread taxa at the soil-root interface. **a**, Correlation between OTUs prevalence across sites in Soil, rhizosphere (RS), rhizoplane (RP), Root and averaged OTUs relative abundance (RA, log2). Bacteria: upper panels. Fungi: middle panels. Oomycetes: lower panels. Blue: geographically restricted OTUs (site prevalence < 20%). Orange: geographically common OTUs (site prevalence 20-80%). Red: geographically widespread OTUs (site prevalence > 80%). For calculating averaged RA, only samples where the actual OTUs are present were considered. The different shapes highlight OTUs detected one year, or across two or three years. RA and prevalence were averaged across the years where one OTU is present. OTUs with relative abundance < 0.1% were excluded from the datasets. **b**, For each microbial group (bacteria, fungi, and oomycetes), Spearman’s rank correlations (p *<* 0.01) were determined between OTUs prevalence in roots of *A. thaliana* and OTUs prevalence in roots of neighboring grasses. The geographically widespread OTUs detected in roots of *A. thaliana* and grasses are indicated with numbers.

**Supplementary Fig. 4:**
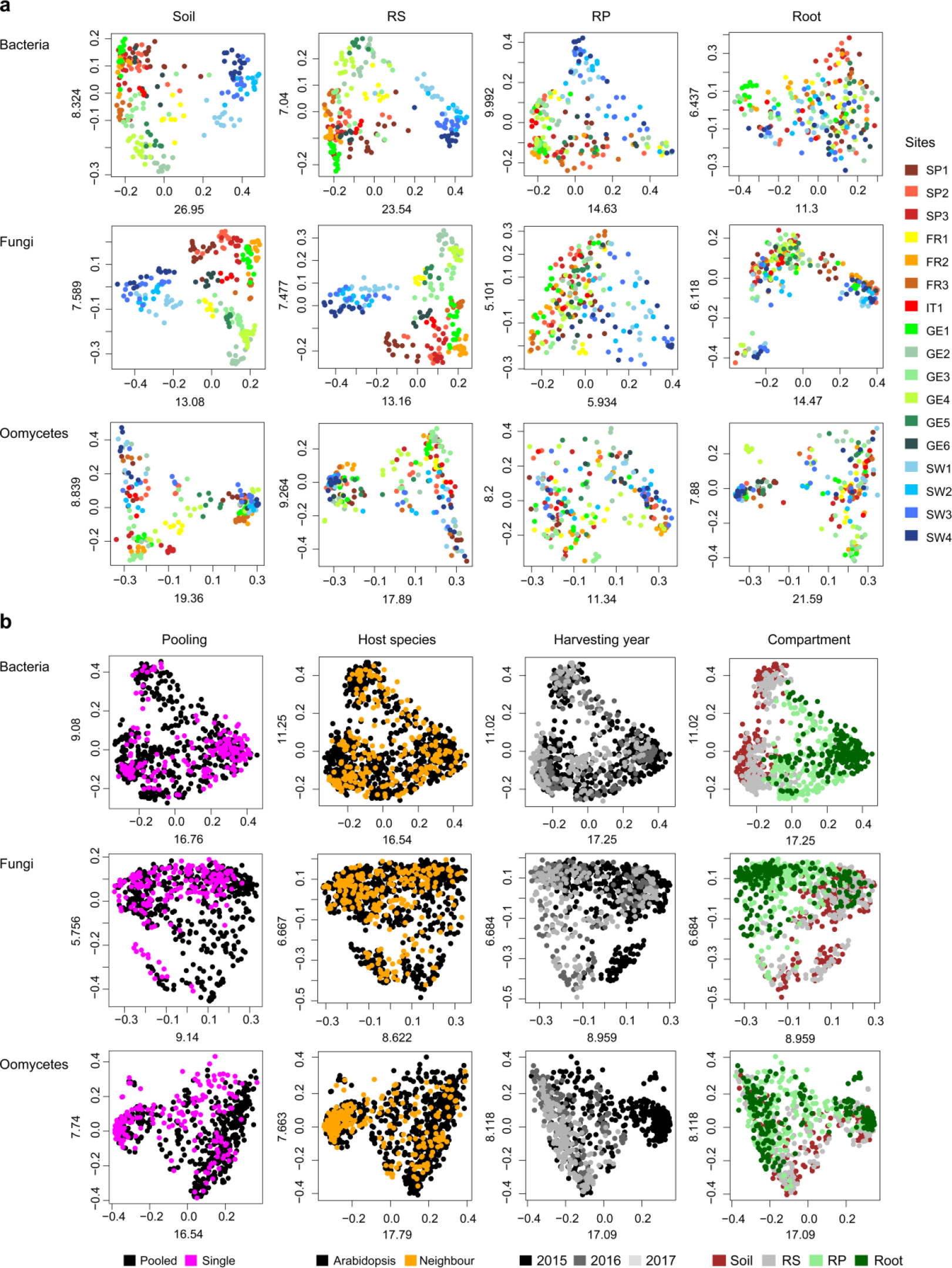
Influence of site, host species, and harvesting year on microbial community structure in *A. thaliana* populations. **a**, Principal coordinate analysis (PCoA) based on Bray Curtis distances for soil-, rhizosphere (RS), rhizoplane-(RP), and root-associated microbial communities detected in 17 sites across three successive years in European *A. thaliana* populations. Microbial communities in each compartment are presented for bacteria (upper panel), fungi (middle panel), and oomycetes (lower panel) and color-coded according to the site. **b**, PCoA based on Bray Curtis distances between samples (bacteria n = 881, fungi n = 803, oomycetes n = 875) harvested across 17 sites, four compartments and three successive years. Microbial communities are presented for the whole dataset for bacteria (upper panel), fungi (middle panel), and oomycetes (lower panel) and color-coded either according to the root pooling strategy (root from a single individual compared to pooled roots from four individuals), the harvesting year (2015, 2016, 2017), or the compartment. Additional soil, RP, RS, and Root samples (bacteria n = 238, fungi n = 241, oomycetes n = 236) from neighboring grasses were included in the PCoA plot where samples are color coded according to the host species. OTUs with relative abundance < 0.1% were excluded from the datasets.

**Supplementary Fig. 5:**
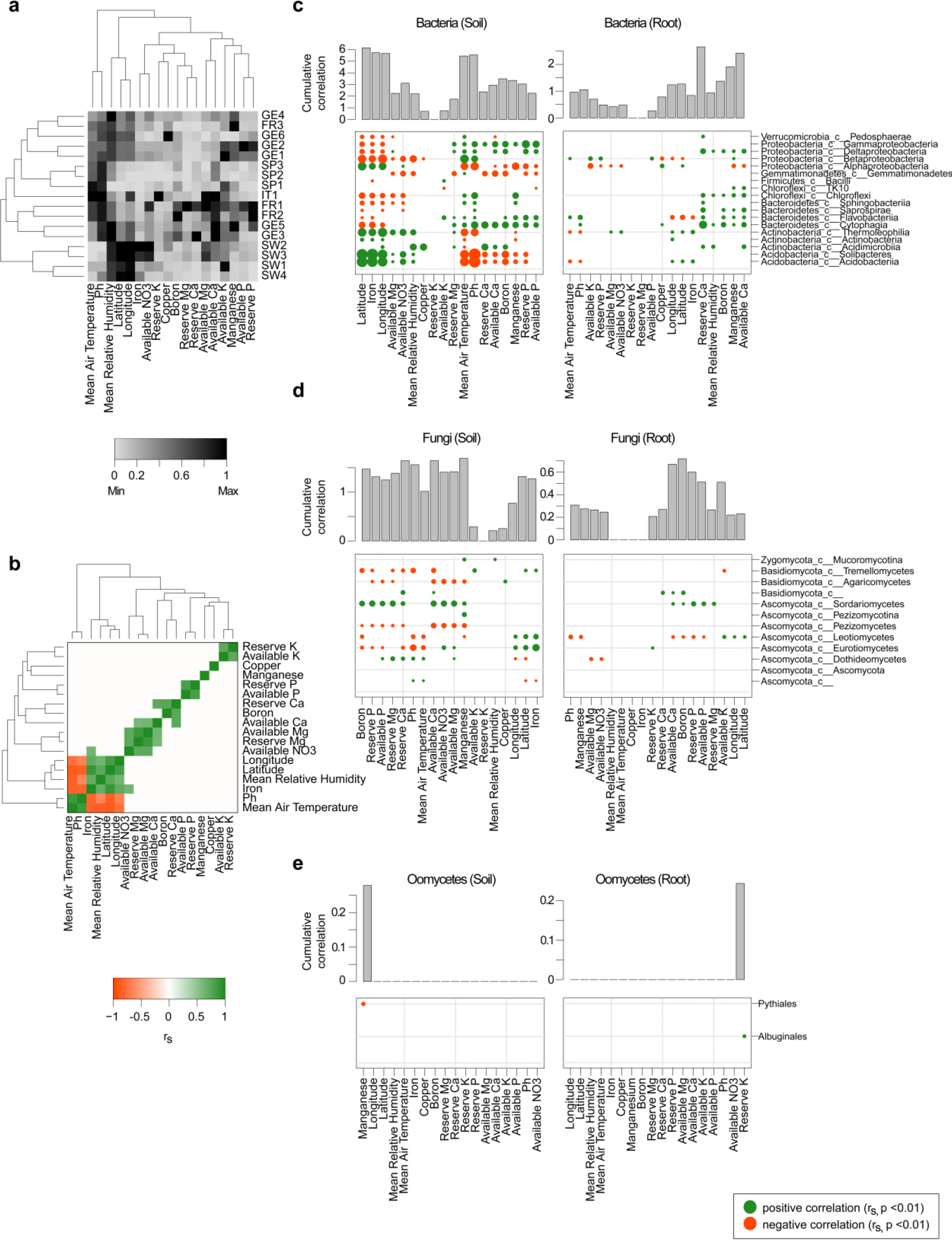
Association between local factors and microbial abundance profiles. **a**, Heatmap showing the distribution of soil properties among each of the 17 sites. Real property values were normalized (0 = lowest measured value, 1 = highest measured value). **b**, Heatmap showing significant correlations detected between properties using data from all sites (Spearman’s rank correlation, p < 0.01). **c**, Correlation between properties and RA of bacterial taxa (aggregated at class level) in soil samples (left panel) and root samples (right panel) (Spearman’s rank correlation, p < 0.01). **d**, Correlation between properties and RA of fungal taxa (aggregated at class level, if available) in soil samples (left panel) and root samples (right panel) (Spearman’s rank correlation, p < 0.01). **e**, Correlation between properties and RA of oomycetal taxa (aggregated at order level, if available) in soil samples (left panel) and root samples (right panel) (Spearman’s rank correlation p < 0.01). Size of circles in **c, d**, and **e** is proportional to measured r values. The respective barplots show the cumulative correlation score for each variable.

**Supplementary Fig. 6:**
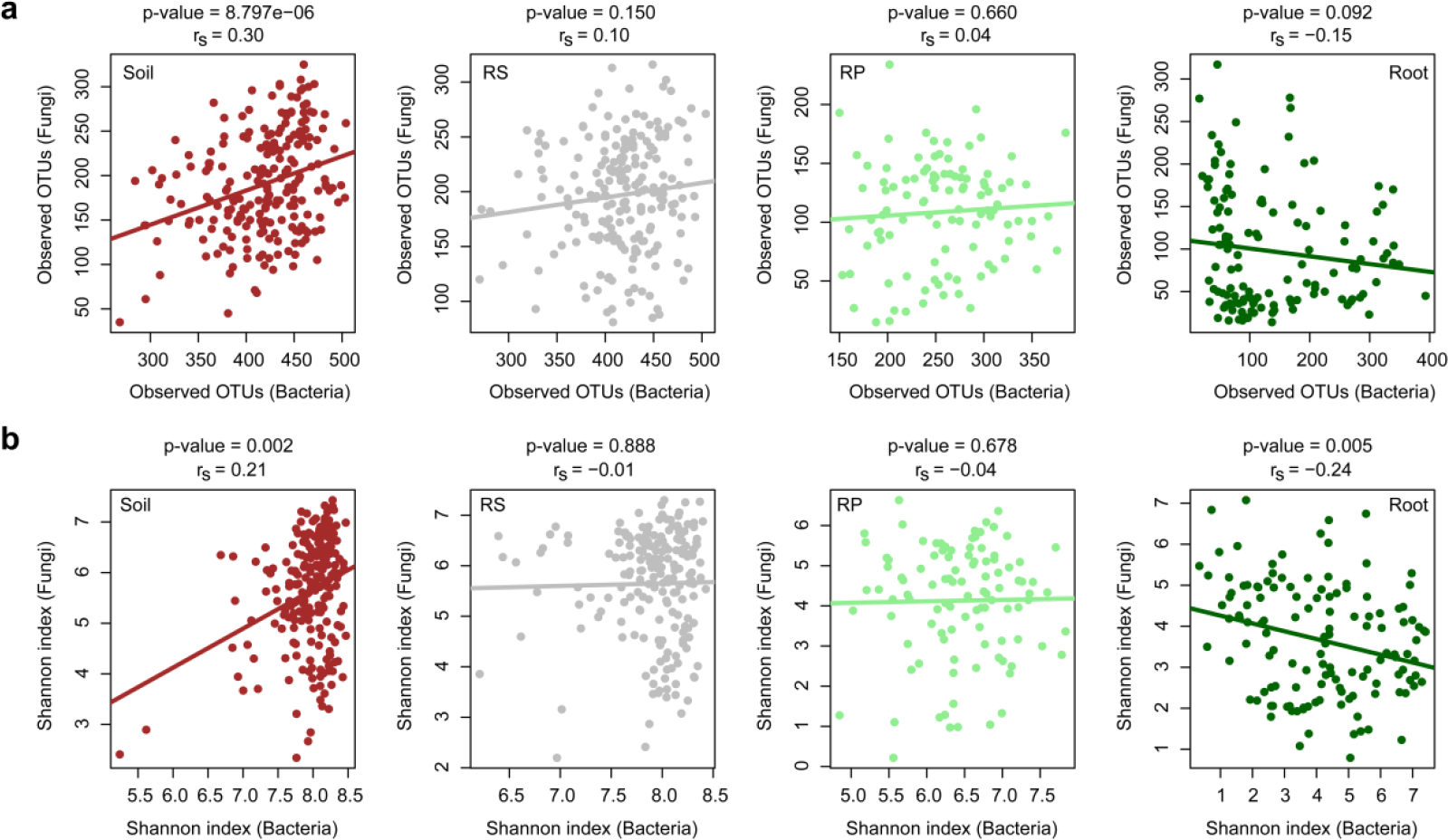
Correlation between bacterial and fungal alpha diversity across compartments. **a**, Spearman’s rank correlation between the number of observed fungal OTUs and the number of observed bacterial OTUs in soil, rhizosphere (RS), rhizoplane (RP), and root samples. All soil, RS, RP, and root samples from the 17 sites were taken into account and the datasets were rarefied to 1,000 reads (soil: n = 212, RS: n = 197, RP: n = 104, Root: n = 131). The Spearman’s rank correlation coefficient and associated p-values are depicted above each graph. **b**, Spearman’s rank correlation between the fungal Shannon index and the bacterial Shannon index in Soil, RS, RP, and root samples. All samples from the 17 sites were taken into account and the datasets were rarefied to 1,000 reads (soil: n = 212, RS: n = 197, RP: n = 104, root: n = 131). The Spearman’s rank correlation coefficient and associated p-values are depicted above each graphs.

**Supplementary Fig. 7:**
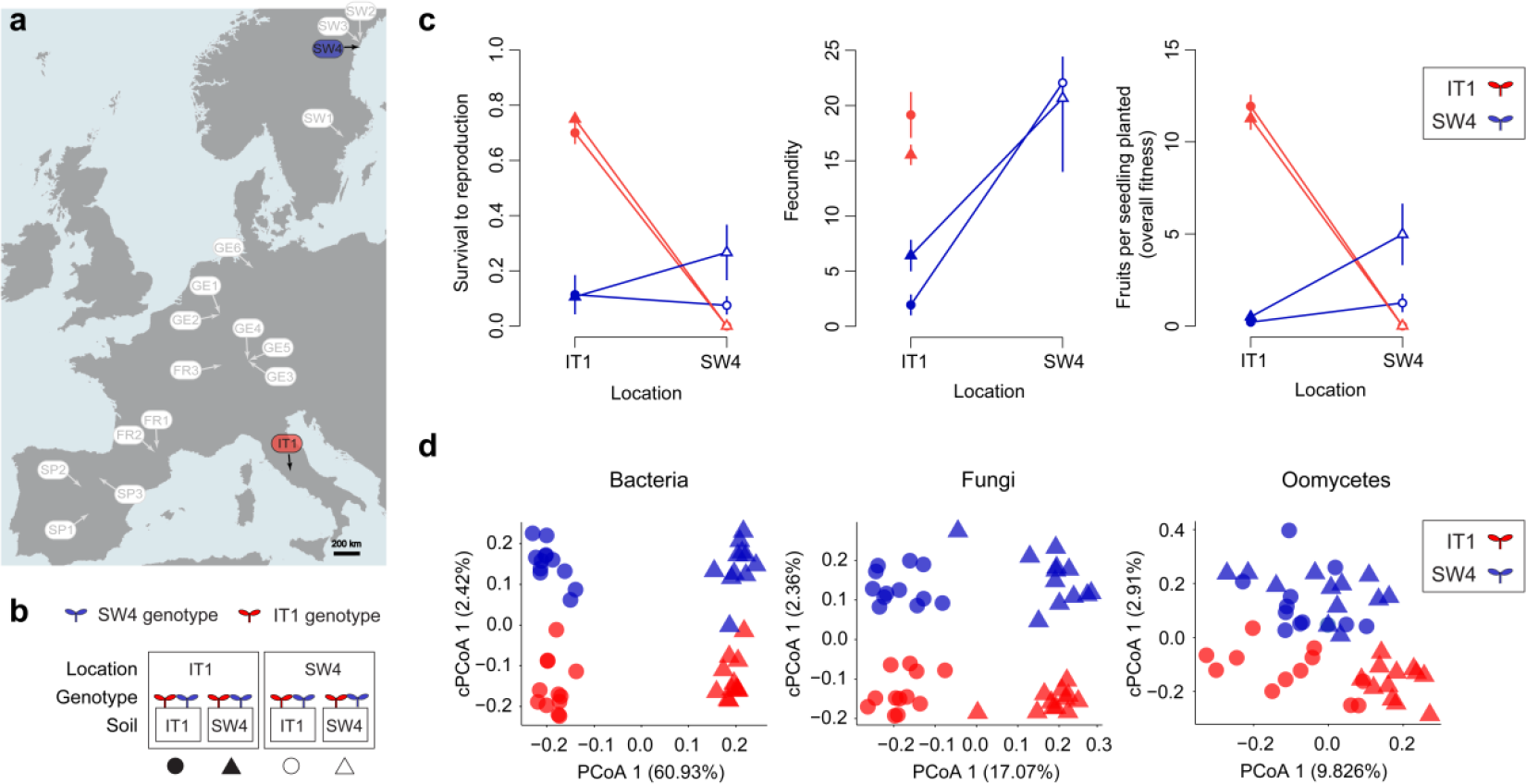
Reciprocal transplant between two *A. thaliana* populations in Sweden and Italy. **a**, European map showing names and locations of the 17 *A. thaliana* populations. The IT1 and SW4 sites selected for the reciprocal transplant experiment are highlighted in red and blue, respectively. **b**, Schematic overview of the reciprocal transplant experiment. Soils and plant genotypes from IT1 and SW4 sites were reciprocally transplanted in the two locations (eight different treatment combinations). The symbols below the schematic view correspond to the symbols used in panels c and d. **c**, Fitness of Italian and Swedish genotypes (red and blue color, respectively) when reciprocally planted in Italian and Swedish soils (circle and triangle symbols, respectively) and grown at Italian and Swedish locations (filled and open symbols, respectively). Plant survival, fecundity (number of fruits per reproducing plant), and overall fitness (number of fruits per seedling planted). Means based on block means ± SE are given. Note that no Italian plant survived to reproduce at the Swedish site. **d**, Bray-Curtis distances constrained by genotype for bacterial, fungal, and oomycetal communities in whole root samples (cPCoA, see axis 2). Results are shown for Italian and Swedish genotypes (red and blue color, respectively) planted in Italian and Swedish soils (circle and triangle symbols, respectively) at IT1 site only since no Italian plant survived at the SW4 site. The percentage of variation explained by the two genotypes is plotted along the second axis and refers to the fraction of the total variance of the data that is explained by the constrained factor (i.e: genotype; bacteria p = 0.001; fungi p = 0.026; oomycetes p = 0.002).

**Supplementary Fig. 8:**
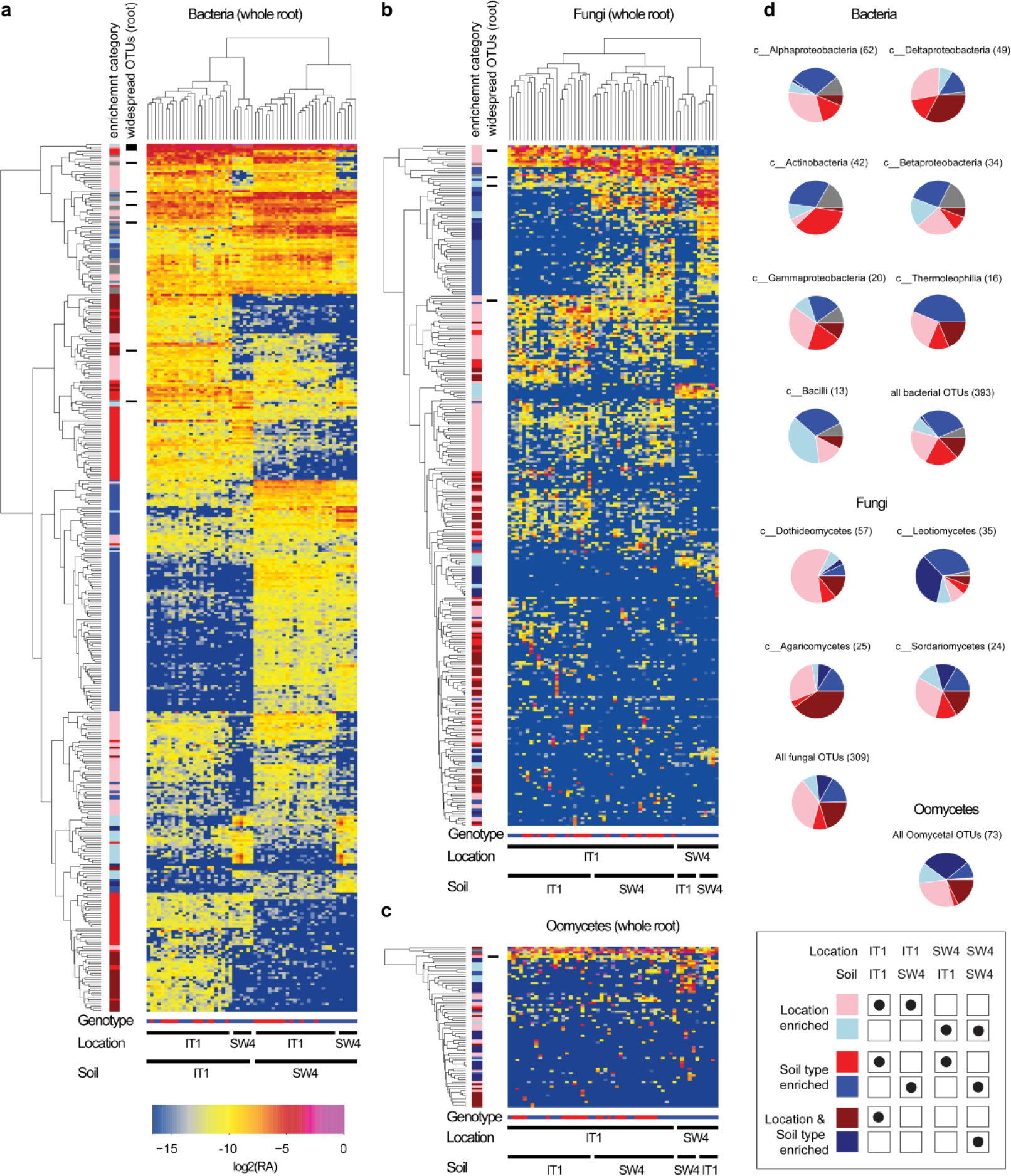
OTU distribution pattern across root samples in the transplant experiment. **a**, Heatmap depicting the relative abundance (log2) of bacterial OTUs in roots of Italian and Swedish genotypes grown in Italian and Swedish soils at IT1 and SW4 locations. OTUs and samples are hierarchical clustered. Enrichment patterns of each OTU was estimated according to the categories described in the lower right side of the figure and highlighted with different colours next to the heatmap. The relative abundance of OTUs falling into one of the six categories is always higher in that category compared to the mean relative abundance measured across all samples. OTUs that are present in all samples (relative abundance > 0.1%) and did not fall in any of the six categories are marked in grey. The heatmap is filtered for OTUs that have at least an average relative abundance of 0.01% across all root samples. Samples have been filtered to contain at least 1,000 reads. Genotype of plants for each sample is indicated below each heatmap. Blue: Swedish genotype. Red: Italian genotype. Note that no Italian plant survived at the Swedish site. **b**, Heatmap depicting the relative abundance (log2) of fungal OTUs in roots of Italian and Swedish genotypes grown in Italian and Swedish soils at IT1 and SW4 locations. **c**, Heatmap depicting the relative abundance (log2) of oomycetal OTUs in roots of Italian and Swedish genotypes grown in Italian and Swedish soils at IT1 and SW4 locations **d**, Percentage of OTUs falling into one of the six categories are presented as pie charts for each main taxonomic classes. The number of OTUs that belong to each microbial class is given in brackets.

## Supplementary Tables

**Supplementary table 1: European sites from where *Arabidopsis thaliana* and grasses populations were harvested.**

**Supplementary table 2: Primers utilized in this study to profile bacterial, fungal and oomycetal communities in soil and root samples.**

**Supplementary Table 3: Microbial communities’ variation explained by several factors across all compartments.**

**Supplementary Table 4: Description of geographically widespread OTUs detected in A. thaliana Root samples.**

**Supplementary Table 5: Microbial communities’ variation explained by several factors and environmental variables for each individual compartment.**

**Supplementary Table 6: Microbial communities’ variation explained by host species at each site in the root compartment.**

**Supplementary table 7: Microbial communities’ variation explained by compartment and by soil, location, and genotype in a field reciprocal transplant experiment**

**Supplementary Table 8. Survival, number of fruits produced by survivors, and the number of fruits per seedling planted in the reciprocal transplant experiment conducted at the sites of the IT1 and SW4 populations.**

**Supplementary Table 9. Analysis of effects of soil (Italian vs. Swedish), and genotype (Italian vs. Swedish) on total fitness (number of fruits per seedling planted) in a field experiment conducted at the site of the Italian genotype.**

